# GpsB coordinates cell division and cell surface decoration by wall teichoic acids in *Staphylococcus aureus*

**DOI:** 10.1101/2021.09.29.462461

**Authors:** Lauren R. Hammond, Sebastian J. Khan, Michael D. Sacco, Catherine Spanoudis, Abigail Hough, Yu Chen, Prahathees J. Eswara

## Abstract

Bacterial cell division is a complex and highly regulated process requiring the coordination of many different proteins. Despite substantial work in model organisms, our understanding of the systems regulating cell division in non-canonical organisms, including critical human pathogens, is far from complete. One such organism is *Staphylococcus aureus*, a spherical bacterium that lacks known cell division regulatory proteins. Recent studies on GpsB, a protein conserved within the Firmicutes phylum, have provided insight into cell division regulation in *S. aureus* and other related organisms. It has been revealed that GpsB coordinates cell division and cell wall synthesis in multiple species by interacting with Penicillin Binding Proteins (PBPs) and other partners. In *S. aureus*, we have previously shown that GpsB directly regulates FtsZ polymerization. In this study, using *Bacillus subtilis* as a tool, we isolated intragenic and extragenic spontaneous suppressor mutants that abrogate the lethality of *S. aureus* GpsB overproduction in *B. subtilis*. Through characterization of these mutants, we identified several key residues important for the function of GpsB. Furthermore, we discovered an additional role for GpsB in wall teichoic acid (WTA) biosynthesis in *S. aureus*. Specifically, we show that GpsB directly interacts with the wall teichoic acid export protein TarG using a bacterial two-hybrid analysis. We also identified a three-residue motif in GpsB that is crucial for this interaction. Based on the analysis of the localization of TagG in *B. subtilis* and its homolog TarG in *S. aureus*, it appears that WTA machinery is a part of the divisome complex. As such, we show additional evidence to the growing body of work that suggests that along with peptidoglycan synthesis, WTA biosynthesis and export may take place at the site of cell division. Taken together, this research illustrates how GpsB performs an essential function in *S. aureus* by directly linking the tightly regulated cell cycle processes of cell division and WTA-mediated cell surface decoration.

**IMPORTANCE/AUTHOR SUMMARY:** Cytokinesis in bacteria involves an intricate orchestration of several key cell division proteins and other factors involved in building a robust cell envelope. One of the key factors that differentiates Gram-positive bacteria from Gram-negative bacteria is the presence of teichoic acids interlaced within the Gram-positive cell wall. By characterizing the role of *Staphylococcus aureus* GpsB, an essential cell division protein in this organism, we have uncovered an additional role for GpsB in wall teichoic acids (WTA) biosynthesis. We show that GpsB directly interacts with TarG of the WTA export complex. We also show this function of GpsB may be conserved in other GpsB homologs as GpsB and the WTA exporter complex follow similar localization patterns. It has been suggested that WTA acts as a molecular signal to control the activity of autolytic enzymes, especially during the separation of conjoined daughter cells. Thus, our results reveal that GpsB, in addition to playing a role in cell division, may also help coordinate WTA biogenesis.

## INTRODUCTION

One of the defining characteristics of life is the ability for a cell to grow and divide. Although there are some exceptions, the predominate process for growth and division in bacteria is binary fission where one bacterial cell grows and divides to produce two similarly sized daughter cells [1, 2]. This tightly regulated process coordinates chromosome segregation, cell elongation, and controls the activity of cell division proteins [3, 4]. Although significant strides have been made to identify the molecular mechanism regulating the cell division machinery, gaps remain in our knowledge, particularly in the non-model organisms [2, 5]. For example, GpsB (the central protein of interest in this study) plays an important, and in some cases essential, role in the cell growth regulation of multiple clinically-relevant Gram-positive organisms (specifically Firmicutes), but is absent in Gram-negative organisms [6, 7]. This highlights the need for a closer analysis of such processes and proteins in multiple species.

In *Bacillus subtilis*, *Listeria monocytogenes* and *Streptococcus pneumoniae*, GpsB links cell wall biosynthesis with the cell division process by interacting with penicillin binding proteins (PBPs), thereby helping to regulate and maintain proper cell shapes [8–14]. Our lab reported that *S. aureus* GpsB directly interacts with FtsZ and affects its polymerization characteristics [15]. In our previous study, we described the lethal phenotype associated with the overproduction of *S. aureus* GpsB (GpsB^SA^) in its Firmicutes relative *B. subtilis*. In this study, we utilized this phenotype to conduct a suppressor screen to identify residues that are important for the function of GpsB^SA^ and pathways through which GpsB^SA^ may exert its function. Herein, we describe the effects of seven intragenic GpsB^SA^ suppressor mutations and characterize their ability to abrogate cell division inhibition. Additionally, we investigated extragenic suppressor mutations through whole genome sequencing that allowed us to delineate a novel role for GpsB in linking central cell division directly to the wall teichoic acids (WTA) pathway. Specifically, suppressor mutations were mapped to *tagG/tagH* genes in *B. subtilis*, and subsequent bacterial two-hybrid (BACTH) analysis confirmed the direct interaction between GpsB^SA^ and the *S. aureus* homolog of TagG, TarG. We also show a 3-amino acid motif, positioned away from the well-characterized PBP binding site, that appears to be important for TarG binding. Furthermore, we show that the interaction between GpsB and the WTA export complex may be conserved beyond *S. aureus*, as we also note that TagH has similar spatiotemporal localization pattern as GpsB in *B. subtilis* cells. In *S. aureus*, treatment with antibiotics that target the WTA pathway drastically alters the localization pattern of GpsB, although targeting of GpsB to new division sites remain unaffected. Thus, it appears that GpsB, an essential protein in *S. aureus*, coordinates cell division and WTA production/transport through direct interaction with FtsZ and TarG respectively.

## RESULTS

### Isolation of suppressor mutations of GpsB^SA^ overproduction in *B. subtilis*

To build upon our previous report and further explore other cell cycle processes that involve GpsB^SA^ in an unbiased manner, we isolated suppressors that can tolerate the lethal overproduction of GpsB^SA^ in *B. subtilis* [15]. Briefly, a *B. subtilis* strain harboring an IPTG-inducible *gpsB*^SA^-*gfp* was streaked out on plates containing IPTG and incubated overnight. Following incubation, the colonies that appeared on the plates containing the inducer were presumed to contain spontaneous suppressor mutations. Non-GFP producing isolates were discarded, as the likely cause of the suppression of lethality could be a promoter mutation turning off the expression of *gpsB^SA^-gfp*, a frameshift mutation, or a premature truncation. After multiple rounds of confirmatory screening, the mutations were classified to be either intragenic or extragenic (**Fig. 1A**) [16]. Using this method, we isolated seven intragenic mutations, Y14F, L35S, D41N, D41G, R72H, as well as a deletion and repeat of a 3-amino acid stretch 66-68 LEE (ΔLEE and LEErpt) that are listed in **Fig. 1B**, of these L35S was reported previously [15]. Throughout the manuscript, these suppressor mutations as a group will be referred to as *GpsB-GFP. We then analyzed the multiple sequence alignment of GpsB from *S. aureus*, *B. subtilis*, *L. monocytogenes*, and *S. pneumoniae* (**Fig. 1C**). Of the first four mutations (Y14F, L35S, D41N, and D41G), the latter three occur in highly conserved residues. Tyr14 is wedged between a conserved Lys (Lys11) which was reported to be important for forming a bidentate salt bridge with the proximal glutamate/aspartate (Glu15 in *S. aureus*) in other organisms [6]. Of note, Phe replaces Tyr in the corresponding position (Tyr14 of *S. aureus*) in *S. pneumoniae* GpsB. The remaining three mutations (ΔLEE, LEErpt, and R72H) are near the disordered linker connecting the N- and C-terminal domains (**Fig. 1C**). The Leu, Glu, Glu (LEE) motif is conserved in *B. subtilis*, however Arg72 is less conserved but appears in *L. monocytogenes*.

**Figure 1.**
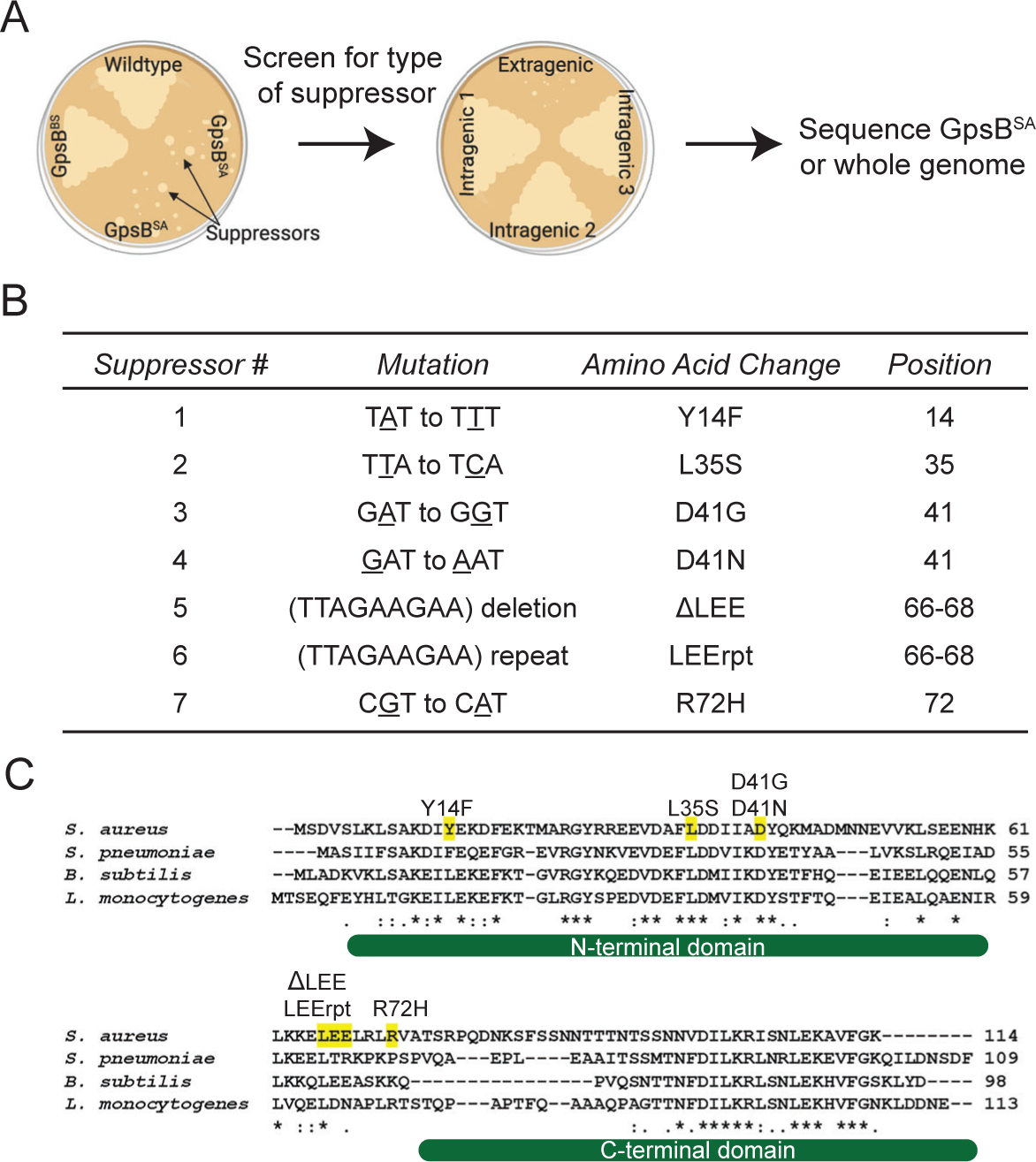
GpsB^SA^-GFP Suppressor Screen. (A) Cartoon diagram depicting workflow for GpsB^SA^-GFP suppressor screen. Figure generated using Biorender.com (B) Table describing seven isolated GpsB^SA^-GFP intragenic suppressor mutations detailing the mutation, amino acid change, and amino acid position. (C) Multiple sequence alignment of GpsB from *S. aureus*, *S. pneumoniae*, *B. subtilis*, and *L. monocytogenes*. Intragenic suppressor mutations identified in the GpsB^SA^-GFP suppressor screen are highlighted in yellow and labeled above the residue locations. *,:, and., indicate fully, strongly, or weakly conserved residues respectively. The structured N- and C-terminal domains are shown under the sequence alignment as reported previously [6].

### Suppressor mutations of GpsB^SA^ abolish cell division inhibition in *B. subtilis* cells

To examine the ability of *B. subtilis* to tolerate the expression of \**gpsB*^SA^-*gfp*, we conducted a spot titer assay with a clean copy of *gpsB^SA^-gfp* harboring the suppressor mutation cloned into PY79 cells under the control of an inducible promoter, similar to how the wildtype *gpsB^SA^-gfp* was constructed. Cultures containing each of the strains were grown, serially diluted, and then plated onto LB agar plates both with and without inducer (**Fig. 2A**). On the minus inducer plate, all strains were able to grow and no growth defects were noted. On the plus inducer plate, the strain containing unmutated GpsB^SA^-GFP showed a severe growth defect consistent with our previous report. In contrast, the growth on the minus and plus inducer plates for each of the suppressor mutations was indistinguishable from each other. Western blot analysis was used to confirm the stable production of each mutant in *B. subtilis* (**Fig. S1A)**. Although most suppressors are stably produced, L35S displayed a distinct cleavage product **(Fig. S1B)** and LEErpt appears to be more stable as it accumulates to a larger extent even in the absence of the inducer. Regardless, all seven suppressor mutations in GpsB^SA^ allow *B. subtilis* cells to grow on solid medium in the presence of inducer.

**Figure 2.**
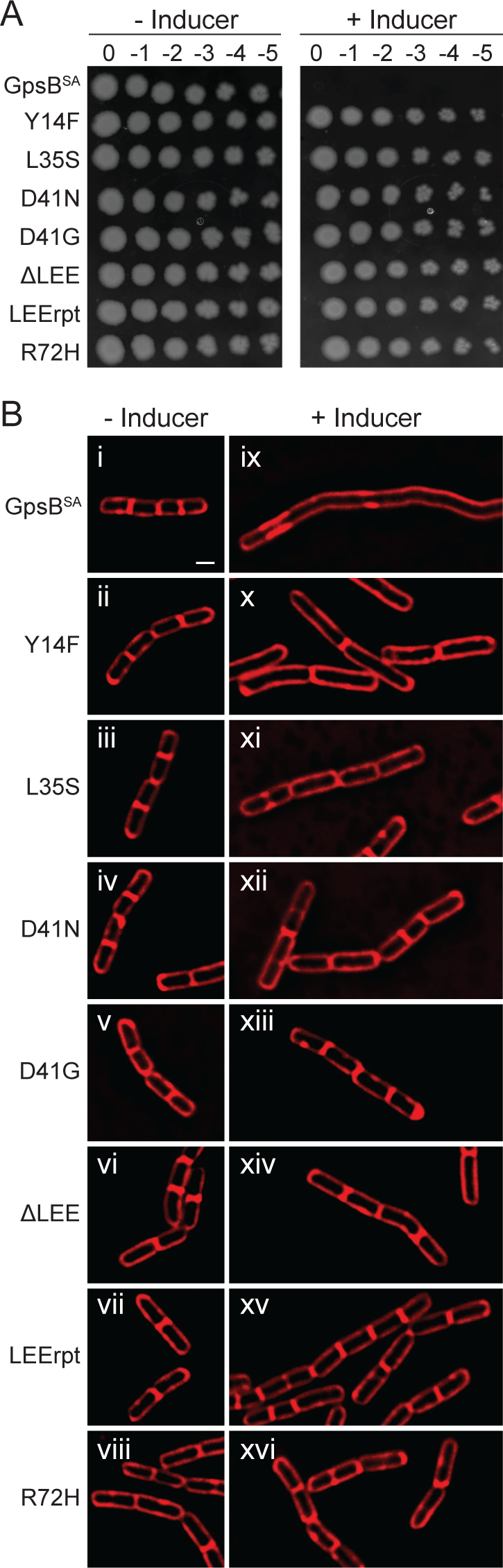
Growth characteristics of intragenic suppressor mutations of GpsB^SA^-GFP. (A) Spot titer assay of unmutated GpsB^SA^-GFP (GG8) (top row) and isolated *GpsB^SA^-GFP intragenic suppressor mutations (CS89-CS93, PE377, and PE448) were serially diluted, spotted on plates, and grown in the absence (left panel) or presence (right panel) of 1 mM IPTG. (B) Cell morphology of cells harboring unmutated GpsB^SA^-GFP (GG8) and the *GpsB^SA^-GFP intragenic suppressor mutations (CS89-CS93, PE377, and PE448) grown in the absence (i-viii) or presence (ix-xvi) of 1 mM IPTG. Images were taken 3 h after addition of inducer. Membrane was stained with SynaptoRed membrane dye. Scale bar is 1 µm.

We previously reported that the growth defect caused by the expression of *gpsB*^SA^ in *B. subtilis* was due to severe filamentation, which is characteristic of cell division inhibition in this organism [15]. To investigate the effect of the suppressor mutations on cell division inhibition, we performed high resolution fluorescence microscopy (**Fig. 2B**).

Upon expression of *gpsB*^SA^-*gfp*, we observed the previously reported cell division inhibition and filamentous phenotype (**Fig. 2B ix**). Notably, upon expression of each of the suppressor mutations, the *B. subtilis* cells no longer display the filamentous phenotype and appear to be dividing normally (**Fig. 2B x - xvi**). We also examined *GpsB-GFP localization in all strains (**Fig. S1C**). Diffused localization was observed for the L35S (as noted previously; [15]) and D41N suppressors (**Fig. S1C iii and v**). Otherwise, all *GpsB-GFP strains displayed a mostly wildtype-like localization pattern.

### Most \**gpsB^SA^* mutants are dominant alleles and can suppress the toxicity of *gpsB^SA^* upon co-expression

To determine if any of the \**gpsB* mutations suppress the toxicity of the wildtype *gpsB^SA^* allele, we engineered a *B. subtili*s strain to co-express both *gpsB^SA^-gfp* and \**gpsB^SA^-gfp* under the control of an IPTG-inducible promoter to produce stoichiometrically equivalent amounts of both wildtype and mutant proteins. We then performed a spot titer assay to examine any growth defects in these strains (**Fig. S2A**). In the strains carrying the suppressor mutations, six were able to restore growth and grow both in the absence and presence of inducer. The seventh mutation, Y14F, showed a weak dominant negative effect as it was able to grow in serial dilutions 2 to 3 log-fold higher than GpsB^SA^-GFP alone. However, it was not as strong as the other mutations that grew in a serial dilution that was 5 to 6 log-fold higher than GpsB^SA^-GFP. Interestingly cells overproducing Y14F variants are on average longer (3.92 µm, n=100) when compared to minus inducer control (2.18 µm, n=100) implying partial functionality of this mutant (compare Fig. 2B panels ii and x). Next, we used a BACTH assay to examine the protein-protein interactions that could explain this dominant negative effect. Since GpsB is known to form a hexamer (trimer of dimers), we tested the ability of each mutant to interact with WT GpsB^SA^ by cloning GpsB^SA^ and *GpsB^SA^ into the BACTH plasmids [17]. Pairs of these plasmids were transformed into BTH101 *Escherichia coli* cells for protein-protein interaction analysis on MacConkey agar and by β cultures (**Fig. S2B**). We found that all the mutants retained their ability to interact with WT GpsB. Thus, we believe that a GpsB^SA^-*GpsB^SA^ interaction is likely the reason for the suppression in toxicity observed in **Fig. S2A**.

As GpsB is an essential protein in *S. aureus*, we wondered if the expression of these dominant suppressor mutation harboring copies of *gpsB* would impair the essential function of GpsB^SA^ in its native organism and be lethal to the cells. This straightforward analysis was complicated by the fact that overproduction of *gpsB-gfp* in *S. aureus* by itself is toxic and results in cell enlargement as reported previously [15]. Thus, we were not able to conduct a thorough analysis. However, among these suppressors it appears that Y14F, ΔLEE, and R72H are the most potent in inhibiting the function of native GpsB, as colony formation is almost completely eliminated upon their overproduction (**Fig. S2C**). Stable production of *GpsB^SA^-GFP in *S. aureus* cells was also confirmed through western blotting (**Fig. S2D**). Despite being one of the most toxic mutations, the ΔLEE mutant accumulates to a much lower extent than the other mutants.

### Structural analysis of GpsB^SA^ suppressor mutations

The N-terminal domain of GpsB is characterized by an elongated coiled-coil dimer that is highly conserved among homologs (**Figs. 3 and S3**) and is highly similar to the lipid-binding domain of DivIVA [6]. The GpsB monomer has two regions of organized α-helix of approximately 35-40 residues (α-helix 2) and a secondary structure: a long α-helix of approximately 8 residues (α-helix 1) (**Fig. 3A**). Near the α-helix 1 from one protomer converges at the interface of α-helix 1 from the other protomer to form a groove that binds to the cytoplasmic N-terminal domain of PBPs. Although the PBP binding site is conserved among GpsB homologs, a conclusive positive interaction between *S. aureus* GpsB and any of the PBPs has not been observed yet. The N-terminal domain of GpsB is connected by a non-conserved, disordered linker to a short, helical C-terminal domain.

**Figure 3.**
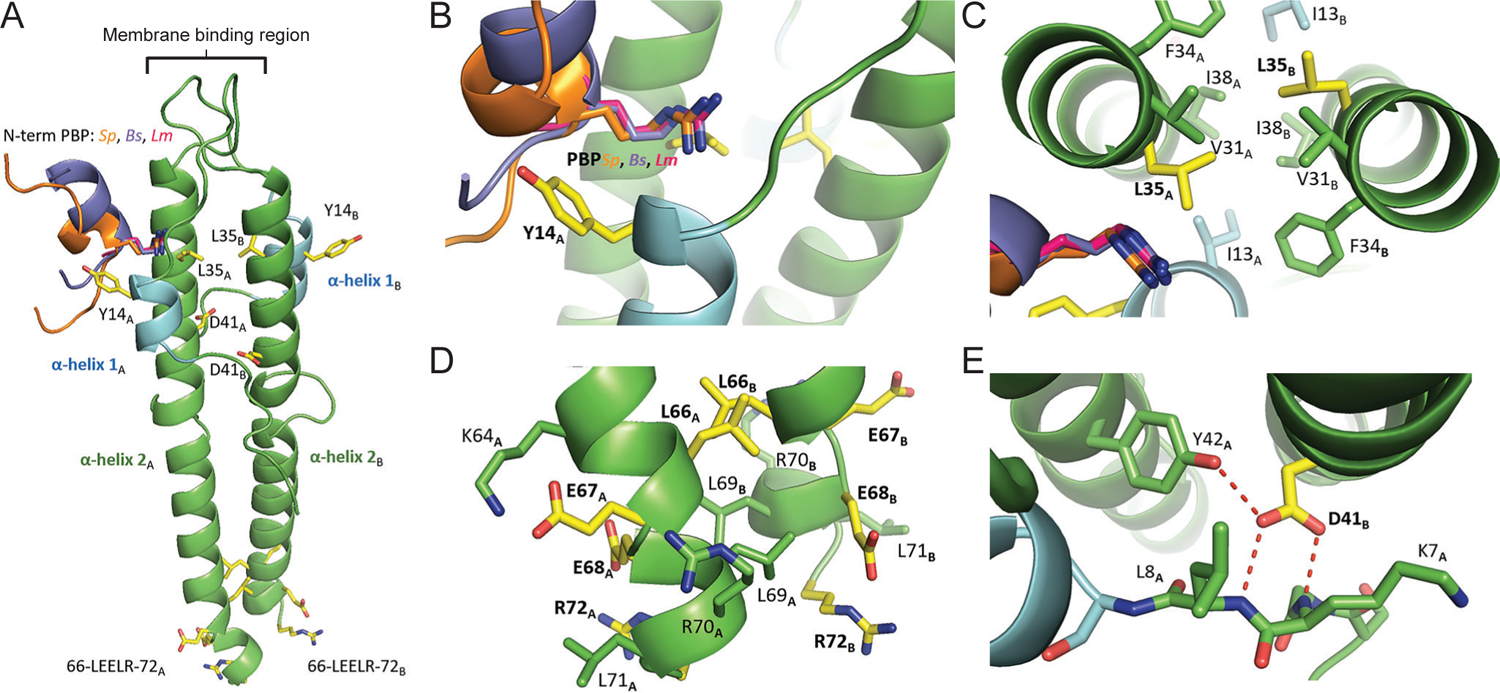
Homology model of S. *aureus* GpsB. Residues mutated in the experiments are colored in yellow. The 8 residues of α-helix 2 S9-E15 are colored in blue. (A)The N-terminal domains of *S. pneumonia* PBP2a (PDB ID 6GQN, orange), *B. subtilis* PBP1a (PDB ID 6GP7, lavender) and *L. monocytogenes* PBPA1 (PDB ID 6GPZ, magenta) bound to GpsB were superimposed and are shown to illustrate the highly conserved PBP-binding α-helix 1 and points directly towards the putative PBP-binding partner. (C) The L35S mutation occurs at the junction of the α-helices 2**_A,B_** and the PBP-binding groove. (D) The D41N, D41G mutations occur at the interface of the loop formed by the first 8 residues α-helix 2 from protomers A and B. (E) The LEELR sequence is the last of α helix 2 before it transitions into a disordered linker sequence that bridges the N-terminal domain to the C-terminal domain.

Using the SWISS-MODEL homology-server, a homology model for the N-terminus of *S. aureus* GpsB was constructed, allowing us to predict the structural effects of these suppressor mutations (**Fig. 3**) [18]. The mutations described in our experiments likely disrupt the structural integrity of GpsB (L35S, D41N/D41G) or alter the recognition elements required for partner binding (Y14F, R72H, LEE repeat/deletion).

### Tyr14 → Phe

α-helix 1 near the PBP-binding groove where it is oriented outwards, away from the core and into the solvent accessible region (**Fig. 3B**). The aliphatic nature of the Tyr side chain is conserved in *S. pneumoniae*, *L. monocytogenes*, and *B. subtilis* where it is Phe, Leu, and Leu respectively. Using the structures of *L. monocytogenes*, *B. subtilis*, and S. *pneumoniae* GpsB complexed with their PBP binding partner (PDB IDs 6GPZ, 6GP7, and 6GQN, respectively), we observe that Tyr14 projects directly towards the PBP N-terminal helix [10, 14]. Because Tyr14 does not interact with other residues in GpsB and the Tyr → Phe mutation is relatively minor, a likely scenario for the more toxic phenotype (**Fig. S2C**), is that Tyr14 facilitates interaction with a binding partner. Furthermore, because the Tyr Phe mutation corresponds to the loss of a phenolic hydroxyl group, the interaction likely involves the formation of a hydrogen bond.

### Leu35 → Ser

Leu35 is positioned at the interface of the PBP binding site and the core of the coiled coil junction (**Fig. 3C**). Despite interacting with adjacent hydrophobic residues near the interfacial core, Leu35 is in immediate proximity to the canonical arginine that is required for PBP binding. However, a mutation to a more polar residue that is capable of hydrogen bonding would seemingly improve this interaction. Therefore, it is most likely that the L35S mutation disrupts the core hydrophobic interactions that are critical for maintaining either the overall structure, or the shape of the PBP binding groove. This may explain the diffused localization (**Fig. S1**). Further supporting this hypothesis is an equivalent mutation at this position in *L. monocytogenes* GpsB, L36A that prevents oligomerization of GpsB, presumably due to the disruption of this hydrophobic core [10].

### Asp41 → Asn, Gly

α-helix 2 where it interacts with a loop formed by the first 10 α-helix 1 (**Figs. 3E, S3C**). Given the proximity of Asp41 to Lys7 and the fact that a mutation to the chemically similar, but neutral Asn produces a non-functional variant, one could mistakenly assume this is a critical electrostatic interaction. However, the side chain of Lys7 is 5.8 Å away from the closest Asp side chain oxygen, well beyond the expected range of favorable electrostatic interactions. Furthermore, Lys7 is not conserved amongst *L. monocytogenes* and *S. pneumoniae*; significantly, *S. pneumoniae* GpsB has an isoleucine at this position (**Fig. S3C**). A closer inspection of this region reveals that Asp41 is an important acceptor of three hydrogen bonds: from the strictly conserved Tyr42 of the adjacent protomer, and the amide nitrogen of the Lys7 and Leu8 main chain. The hydrogen bonds with the backbone nitrogen of Lys7 and Leu8 are important interactions because this attracts the α-helix 1 to interact with correctly forming the PBP binding groove. Therefore, the replacement of any of the Asp oxygen atoms, even with a nitrogen hydrogen bond donor, would likely prohibit the formation of these three highly coordinated hydrogen bonds.

### LEE deletion/repeat and Arg72 → His

The last two turns of the GpsB N-terminal α LEELRLR-72. They are followed by a flexible linker region of approximately 20 amino acids that connects to the C-terminal domain. Interestingly, the LEELRLR region is not conserved amongst other Firmicutes and is unique to *S. aureus*. Multiple i +3 and i +4 electrostatic interactions are formed laterally along LEELRLR by Arg and Glu sidechains and Lys64 (**Figs. 3D, S4**). Additionally, the Leu residues interact through hydrophobic interactions in the core with the corresponding residue of the adjacent protomer and the neighboring Leu of its own chain. The insertion or deletion of a LEE sequence would disrupt the complementarity of sidechain interactions and cause charge-charge repulsion. Deletion of LEE eliminates a pair of (N+3/N+4) +/- interactions (K64-E67, E68) and (E68-R72) while adding one pair of +/+ interactions (K63, K64 -E67R) (**Fig. S4C**). A LEE repeat eliminates one +/- interaction (E68-R72) and adds two pairs of -/- interactions: (E67-R70E, L71E) and (E68-L71E, R72E) (**Fig. S4B**). Therefore, the insertion or deletion of LEE will decrease the helical propensity of this region. Because the disruption of secondary structure is restricted to a small region that is adjacent to a disordered linker, the impact on the overall structure of GpsB structure could be minimal, meaning this specific area may have functional importance for binding to other proteins. Possible proteins include FtsZ [15] or other unique interaction partners (such as TarG discussed later in this report) that could interact with GpsB through LEELRLR.

Additionally, Arg72 is either the last residue of α disordered, linker region that connects the N-terminal domain to the C-terminal domain. Either way, it is unlikely to affect the overall structure of GpsB and could also be a critical residue that interacts with another protein, likely through electrostatic interactions with a Glu or Asp residue.

### Isolation of extragenic suppressors reveals a link between GpsB and wall teichoic acid machinery

In addition to the intragenic suppressors described above, we also isolated extragenic suppressors and then analyzed these mutants through whole genome sequencing (**Fig. 1A**). Through this process we isolated three different suppressor mutations independently. Two of the mutants had the same mutation in *tagH* (Y233C) and the third suppressor had a mutation in *tagG* (R20K) (**Fig. 4A**). TagG and TagH work together to form a complex that exports wall teichoic acids (WTA) that are made intracellularly so they can be anchored to the cell wall [19]. As these WTA genes are essential in *B. subtilis*, to confirm that the extragenic suppressors harbor true suppressor mutations we utilized the previously developed essential gene knockdown tool based on CRISPR interference (CRISPRi) with deactivated Cas9 [20], to disrupt the expression of either *tagG* or *tagH*. Briefly, we investigated the fate of wildtype GpsB^SA^-GFP overproducing cells when *tagG* or *tagH* expression was knocked down (+ xylose) or not (**Fig. 4C**). When we imaged these *tagG* or *tagH* strains without xylose (**Fig. 4B i and iii)**, the strains appeared similar to wildtype. Upon addition of xylose, we noted areas of membrane enrichment near cell poles (**Fig. 4B ii and iv)**, consistent with the previous report of bulging [20]. As shown before (**Fig. 2B)**, induction of *gpsB^SA^-gfp* expression with IPTG leads to filamentation (**Fig. 4C i and ii,**). In the CRISPRi strains of *tagG* or *tagH* with GpsB^SA^-GFP, the IPTG-mediated filamentation is also seen in the absence of xylose (no interference in the expression of *tagG* or *tagH*) (**Fig. 4C iv and viii**). Finally, when we add both IPTG and xylose to induce GpsB^SA^-GFP production and knockdown of *tagG* or *tagH*, the cells are no longer filamentous (**Fig. 4C vi and x**), thus confirming that *tagG* and *tagH* are true suppressors of GpsB^SA^-GFP mediated cell division inhibition. Therefore, it is likely that the extragenic suppressor mutations result in dysregulation of the TagGH complex to suppress the lethal overexpression of *gpsB^SA^*.

**Figure 4.**
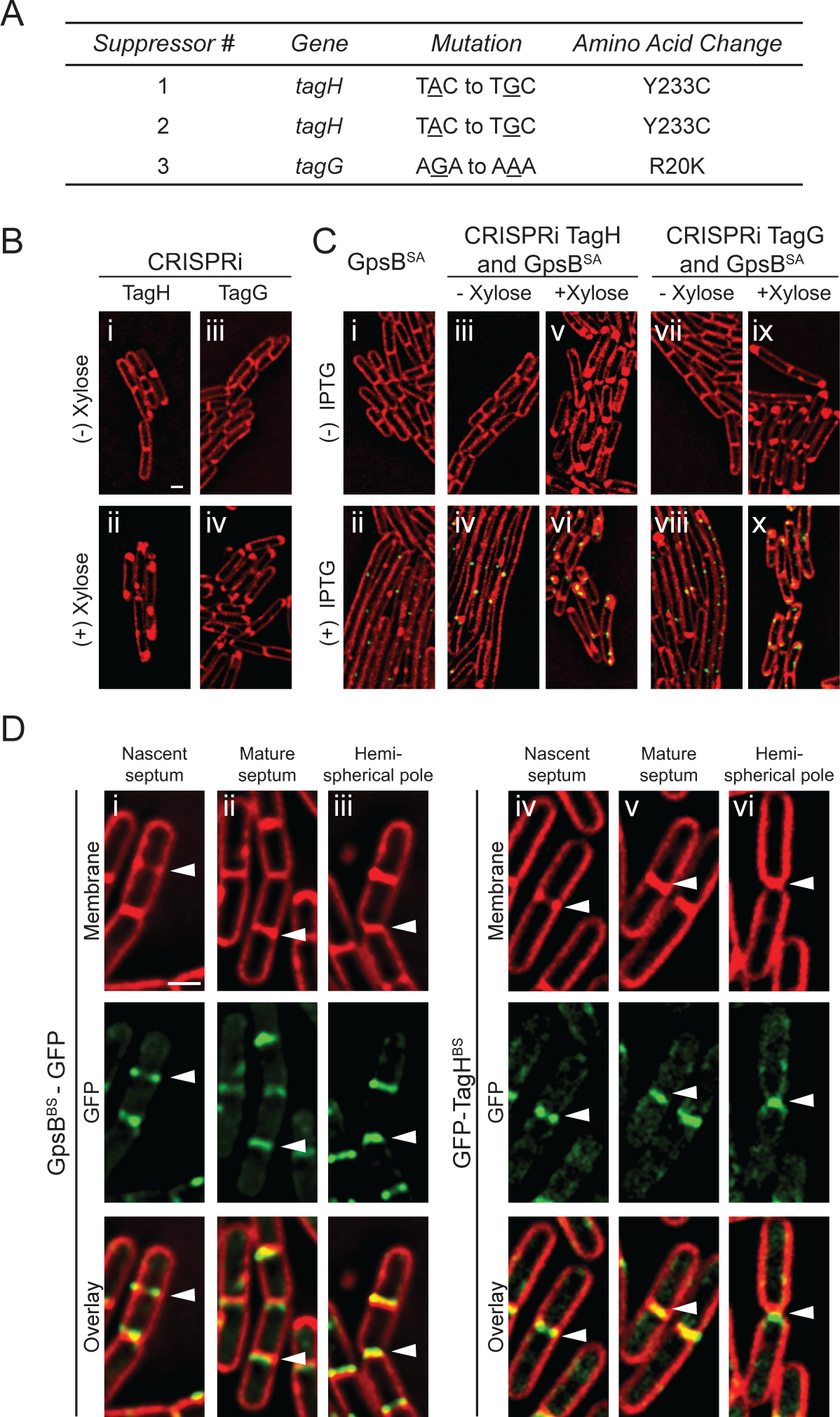
Analysis of GpsB^SA^-GFP extragenic suppressor mutations. (A) Table describing the three isolated extragenic suppressor mutations detailing the specified gene, mutation, and amino acid change. (B) Fluorescence microscopy showing the cell morphology of strains containing a CRISPRi knockdown of *tagH* (SK16) and *tagG* (SK15) in *B. subtilis* grown in the absence (i and iii) and presence (ii and iv) of 1% xylose. (C) Cells containing inducible *gpsB*^SA^-*gfp* (GG8) grown in the absence (i) and presence (ii) of 1 mM IPTG. Strains constructed to have both the CRISPRi knockdown of either *tagH* or *tagG* as well as inducible *gpsB*^SA^-*gfp* (SK18 and SK17) were imaged in the absence of xylose and IPTG (iii and vii), in the presence of xylose only (v and ix), in the presence IPTG only (iv and viii), and finally in the presence of both xylose and IPTG (vi and x). Cells imaged 3 h after the addition of xylose and/or IPTG. Scale bar 1 µm. (D) Fluorescence micrographs tracking the localization pattern of GpsB^BS^-GFP (i – iii) and GFP-TagH^BS^ (PE528) (iv – vi) through different stages of the cell division (see arrowheads). Cell membrane is visualized with SynaptoRed membrane dye. Scale bar 1µm.

To our knowledge, a direct relationship of GpsB and WTA synthesis has not been reported in *B. subtilis*. So, to investigate how TagGH could suppress GpsB^SA^-GFP mediated cell division inhibition, we monitored both GpsB and TagGH localization in *B. subtilis* cells. Using GpsB^BS^-GFP and GFP-TagH^BS^ [21], we analyzed when TagH arrived at the division site (**Fig. 4D**). We found that both GpsB^BS^ and TagH^BS^ arrives at mid-cell early in the division cycle, (at a similar time to GpsB^SA^ in *B. subtilis* [15]), in areas of membrane enrichment (indicating the regions of septal membrane invagination) (**Fig. 4D i and iv; see arrowhead**) and stays at the mature septum (**Fig. 4D ii and v**), at least until the septum transforms into hemispherical cell poles (**Fig. 4D iii and vi)**. This is consistent with the previous reports of TagH^BS^ [21] and GpsB^BS^ localization [8, 9]. Thus, it appears that GpsB may play a role in WTA biosynthesis by interacting with one or more of the WTA biosynthesis proteins, and the toxicity stemming from GpsB^SA^ production in *B. subtilis* could be due to an interaction between GpsB^SA^ and the TagGH^BS^ WTA exporter complex.

### GpsB^SA^ directly interacts with *S. aureus* wall teichoic acid export protein TarG

Next, we analyzed whether GpsB^SA^ could interact directly with the *S. aureus* homologs of TagGH, TarGH (Ta<u>g</u> - teichoic acid <u>g</u>lycerol | Tar - teichoic acid ribitol; [19]), using a BACTH assay. We detected a strong positive interaction between TarG^SA^ and GpsB^SA^ (**Fig. 5A**). We further confirmed the strong positive interaction by quantifying the β-galactosidase enzyme **(Fig S5A)**. Interestingly, a previous work has showed an interaction between GpsB^SA^ and TarO^SA^ via BACTH [22]. Since we could detect an interaction between GpsB and the WTA export complex, we were curious if overexpression of *gpsB* would affect WTA levels in *S. aureus* cells.

**Figure 5.**
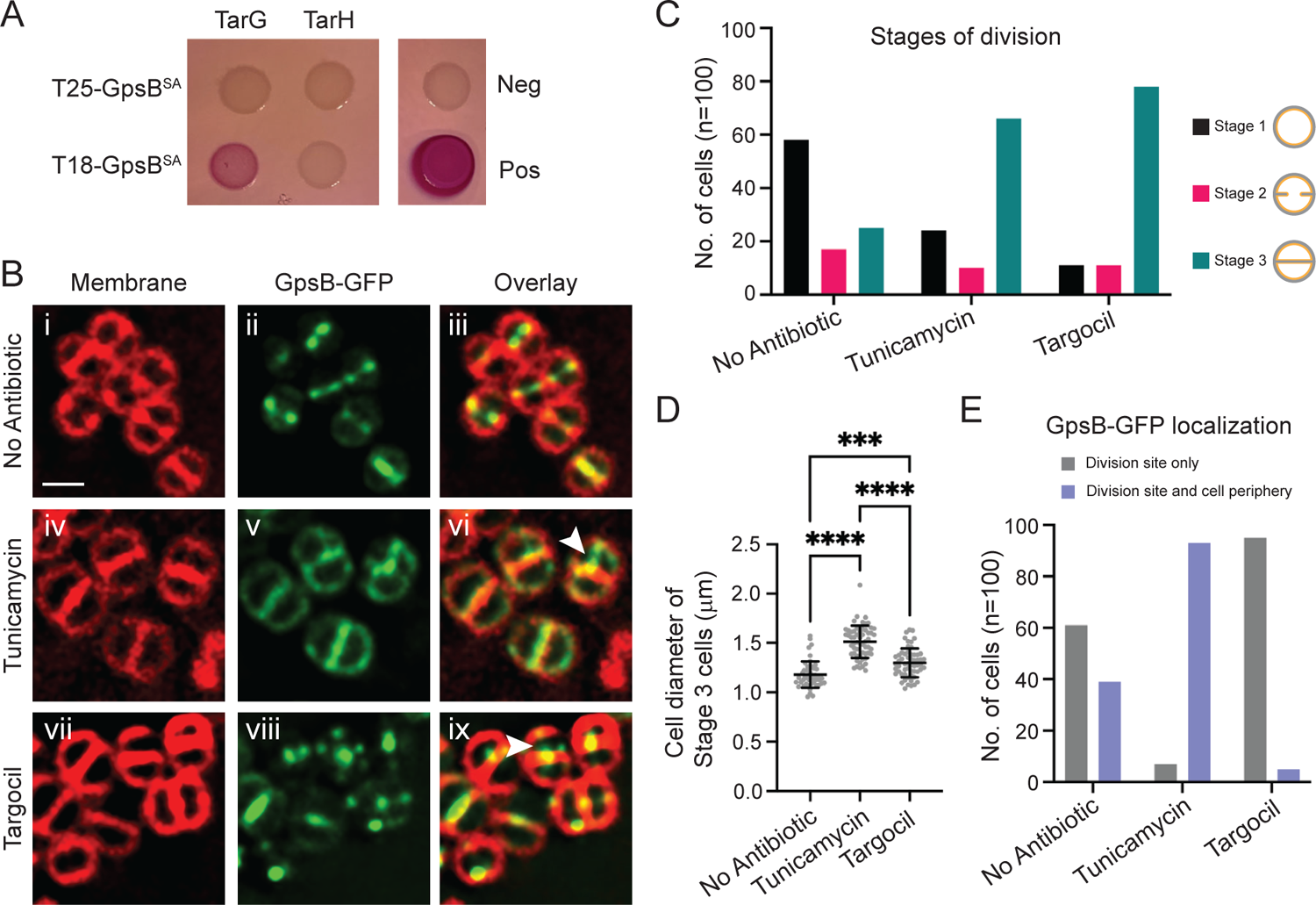
Characterization of the link between WTA exporter TarGH and GpsB in *S. aureus*. (A) Bacterial Two Hybrid Assay to investigate pairwise interactions of GpsB^SA^ with *S. aureus* TarG (SKB1 and SKB2) and TarH (SKB3 and SKB4). A color change to deep pink indicates a positive interaction. Images taken after 24 h of incubation. (B) Localization pattern of GpsB^SA^-GFP (PES6) in SH1000 *S. aureus* cells following 1 h of μg/mL targocil (vii-ix). White arrowheads show instances of GpsB^SA^-GFP localizing to the next division site despite incomplete cell separation. Cell membranes visualized with SynaptoRed membrane dye. Scale bar is 1 µm. (C) Number of cells in each stage of division following 1 h of treatment with tunicamycin or targocil. Stage 1 (black bars) cells show no membrane enrichment at mid-cell. Stage 2 cells (pink bars) have some membrane enrichment at mid-cell but do not have fully formed septa. Stage 3 (teal bars) are cells that have fully formed septa at mid-cell. (D) Quantification of cell diameters from Stage 3 cells treated with tunicamycin or targocil, n = 100 cells, *** P = 0.0002, **** P < 0.0001. One-way ANOVA analysis and multiple comparisons performed in GraphPad Prism 9. (E) Number of Stage 3 cells showing localization of GpsB-GFP at the division site only (grey) or at the division site and the periphery (purple) following 1 h of treatment of tunicamycin or targocil n = 100 cells.

Crude WTA extracts from *S. aureus* cells containing either an empty vector or one overproducing GpsB were visualized under Alcian blue silver staining, but we did not detect any changes in WTA levels in these cells at the condition tested (**Fig. S5B**). In addition, we also observed early division site localization of GFP-TarG (**Fig. S5C**), which supports the physiological significance of GpsB-TarG interaction given that GpsB also localizes to nascent division sites (**Fig. 5B**; [15]). Division site localization of the WTA biosynthesis protein TarO (TagO) has also been noted in *S. aureus* [23]. The division site localization of TarO and TarG and their direct interaction between GpsB reveal that WTA machinery may be part of the divisome complex in *S. aureus*.

### GpsB regulation of FtsZ is independent of wall teichoic acid synthesis

Our previous report showed that GpsB interacts with FtsZ and localizes to the site of division in *S. aureus*, so we wanted to see if GpsB localization at mid-cell was dependent on ongoing wall teichoic acid synthesis/export (**Fig. 5B**). For this purpose, we used two inhibitors of wall teichoic acid synthesis, tunicamycin (early WTA biosynthesis by TarO is inhibited [24]) and targocil (TarGH-mediated WTA export is inhibited [25]). It is noteworthy that in addition to targeting TarO, tunicamycin could also target the early-stage peptidoglycan biosynthesis protein MraY at higher concentration [24]. Based on previous observations, treatment with these antibiotics does not halt cell division, however, the placement of the septa and overall regulation of division is disrupted. Treated cells also had significant cell separation defects presumably due to limited autolysin activity [24–28]. Given this information, we used high resolution fluorescence microscopy to monitor the localization of GpsB^SA^-GFP in cells treated with tunicamycin or targocil. *S. aureus* cells containing GpsB^SA^-GFP were grown to mid-log and then the inducer (IPTG) and the antibiotics (tunicamycin or targocil) were added, and cells were grown for an additional hour. At this point the majority of cells were expected to have completed a full round of division in our experimental condition. In the cultures treated with either tunicamycin or targocil, cells were significantly larger (**Figs. 5B and 5D**). The impaired autolytic activity is also evident with (25%) of cells having a completed septum designated “Stage 3” in the no antibiotic treatment control, vs (66%) in the tunicamycin treated cells and (78%) in the targocil treated cells (**Fig. 5C**). Despite the impaired cell separation in both the tunicamycin and targocil treated cells, we noted evidence of GpsB localizing to sites perpendicular to the previous plane of division (**Fig. 5B vi and ix**; white arrow heads) suggesting that although the cells are not separating properly, they are still attempting to undergo another round of division and that GpsB and presumably FtsZ localization/regulation remain intact. Interestingly, we did note one distinct phenotype between the tunicamycin and targocil treated cells. In the cells treated with tunicamycin, GpsB-GFP appears to remain localized in the peripheral membrane in addition to sites of division, however in the targocil treated cells there is very little peripheral membrane localization (**Figs. 5B** panels **v and viii; and Fig. 5E**).

### Intragenic mutants reveal critical residues for GpsB/TarG interaction

The homology modeling of our intragenic GpsB mutants revealed that three of the ΔLEE, LEErpt, and R72H could disrupt protein-protein interactions. To investigate whether any of these three mutations affected their localization, we imaged these strains under fluorescence microscopy. As longer incubation with ΔR72H is lethal (**Fig. S2C**), cells were imaged at an earlier time point (1 h post-induction). We noted that all three mutations, Δ wildtype-like localization (**Fig. 6A).** Next, we tested the ability of these mutants to interact with TarG. Using the BACTH assay, we tested the interactions of Δ LEErpt, and R72H with TarG. We observed that LEErpt and R72H were still able to ΔLEE no longer interacted with TarG (**Fig. 6B**). This suggests that the presence of these three residues (or the length of the disordered linker connecting the N- and C-terminal domains) are important for the GpsB-TarG interaction. As such, it is tempting to speculate whether the lethality of dominant Δ overproduction (**Fig. S2C**) could be due to the impairment of the native GpsB and TarG interaction.

**Figure 6.**
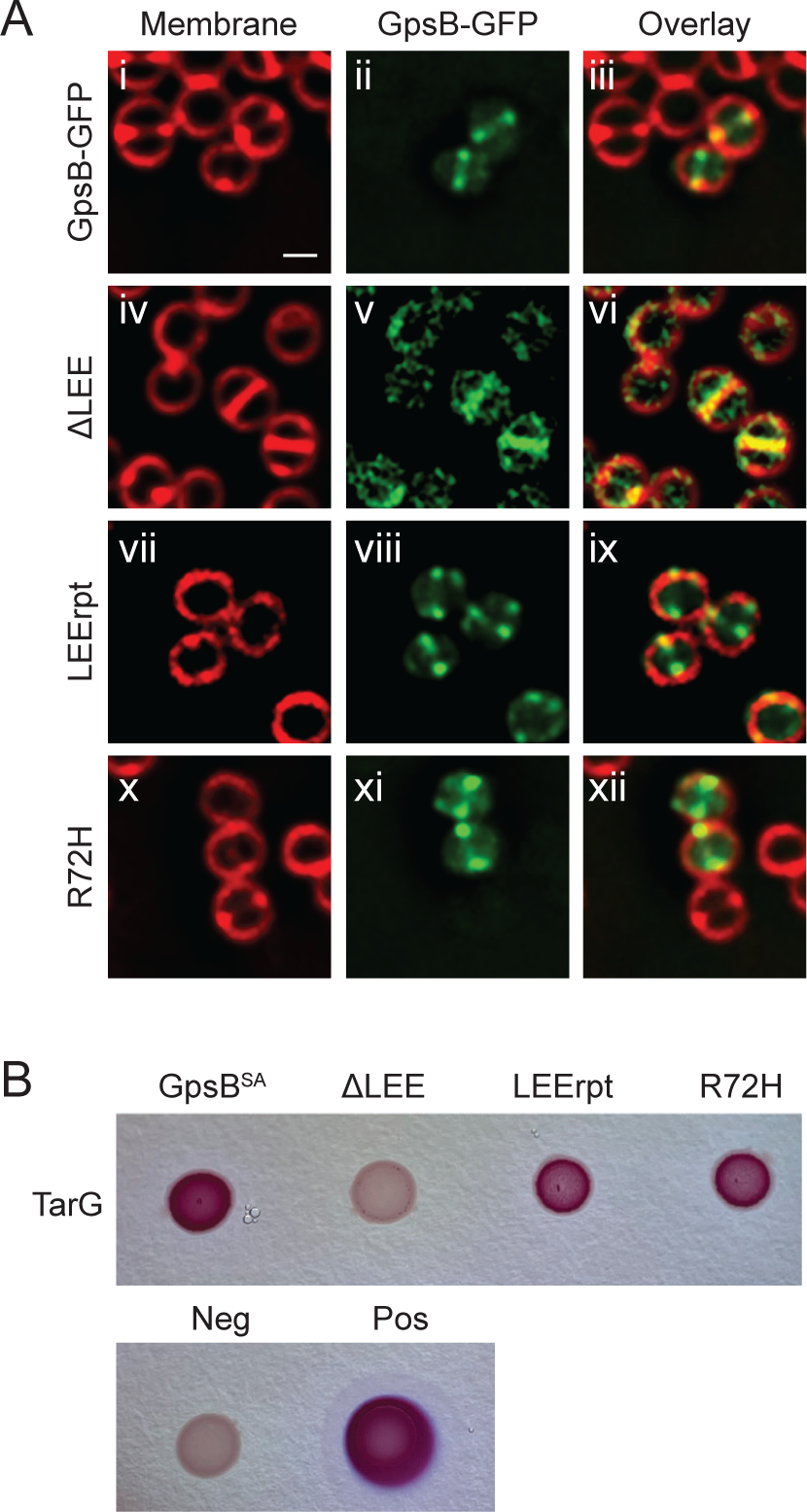
Characterizing GpsB residues important for TarG interaction. (A) Localization pattern of GpsB-GFP (GGS2) or *GpsB-GFP (ΔLEE, LEErpt, R72H) (LH17, LH35, LH18) in RN4220 *S. aureus* cells. Cells imaged 1 h after the addition of 1 mM IPTG and stained with SynaptoRed membrane dye. Scale bar is 1 µm. (B) Bacterial Two Hybrid Assay of pairwise interactions of TarG (SKB2) and GpsB (LH39) or *GpsB (ΔLEE, LEErpt, R72H) (LH47, LH43, LH55). A color change to deep pink indicates a positive interaction. Images taken after 24 h of incubation.

It has been reported that targocil treatment (inhibition of WTA export) could prevent the translocation of major autolysin, Atl [28]. It was shown that *S. aureus* cells treated with targocil exhibited reduced autolysis in comparison to untreated cells. We were interested in testing whether GpsB promoted or inhibited WTA export-mediated autolysis (**Fig. S5D**). The rate of autolysis did not change significantly between cells harboring an inducible copy of *gpsB*, *gpsB*^Δ^, and the empty vector (EV) control.

However, as reported previously, subsequent to targocil treatment, the strain harboring EV control displayed reduced autolysis. Interestingly, in targocil-treated cells overproducing GpsB, the autolysis was reproducibly higher compared to the EV control. This increased autolysis is dependent on TarG interaction, as overproduction of Δ (which lacks the interaction with TarG; **Fig. 6B**) mimicked EV control. Thus, it appears that GpsB may facilitate autolysin (Atl) translocation through its interaction with the TarG component of the WTA export machinery.

## DISCUSSION

GpsB is highly conserved across the Firmicute phylum. In *B. subtilis*, *S. pneumoniae*, and *L. monocytogenes*, GpsB is involved in the coordination of cell division and cell wall synthesis through the binding of PBPs and other partners [6, 7, 14]. Our lab recently reported a novel function of *S. aureus* GpsB in regulating FtsZ polymerization by promoting lateral interactions and facilitating GTP hydrolysis [15]. In this study, we characterized several different mutations in GpsB^SA^ in an effort to more thoroughly understand GpsB mediated cell division regulation in *S. aureus*. Subsequent to a suppressor screen, we identified TarG as a protein interaction partner of GpsB in *S. aureus* and found that residues 66-68 (LEE) of GpsB are critical for this interaction. An earlier BACTH investigation of *S. aureus* cell division factors and peptidoglycan synthesis machinery identified an interaction between EzrA and GpsB, however it failed to show an interaction between GpsB and any other proteins including PBPs (PBP1, PBP2, and PBP3) [29]. However, it appears that GpsB may interact with PBP4 [22], which plays a role in peripheral peptidoglycan synthesis in *S. aureus* [30]. An extensive BACTH-based interaction analysis between lipoteichoic acid (LTA) biosynthesis proteins and cell division factors also revealed that GpsB is not part of the LTA complex in *S. aureus* [31]. Thus, our finding of TarG-GpsB interaction is insightful and was only suspected based on our unbiased approach. In addition to the GpsB interaction with TarG shown here, previous work showed that GpsB interacts with TarO in *S. aureus* (among other proteins) [22].

Traditionally, WTA export has been alluded to occur along the lateral cell wall with the possible help of shape-determining MreB, MreC, and/or MreD proteins in *B. subtilis* [21, 32–34], which could allow for higher order accumulation of WTA at lateral cell wall.

Consistent with this model, several WTA biosynthesis proteins including TagGH also interacts with FtsEX which is involved in cell elongation in *B. subtilis* [35, 36]. However, evidence suggests that WTA synthesis is not only important for cell elongation as a role for FtsEX in cell division and cell separation has been elucidated in *S. pneumoniae* [37] (and it is also known that FtsEX can directly interact with FtsZ in the Gram-negative organism *E. coli* [38]). A role for FtsEX in the activation of a specific peptidoglycan hydrolase has been well documented in these organisms [36–39]. Additionally, of the proteins discussed above, MreC is a known interaction partner of GpsB in *B. subtilis*, *S. pneumoniae*, and *L. monocytogenes* [8, 14]. Interestingly, MreC and other GpsB interaction partners, PBP1 and EzrA also interact with FtsEX in *B. subtilis* [35]. Thus, it is possible to envision a complex made of Mre proteins, FtsEX, PBPs, and GpsB moderating the WTA biosynthesis in *B. subtilis*. Although well-studied MreB is absent in *S. aureus*, MreC and MreD proteins are present and are targeted to the division sites [40], similar to GpsB [15]. Surprisingly, *S. aureus* lack the genes for FtsE and FtsX (L. Aravind - National Center for Biotechnology Information/National Institutes of Health; personal communication). During the course of evolution into a spherical organism from a rod-shaped ancestor, it appears that *S. aureus* has lost the need for the FtsEX complex. However, other alternate mechanism(s) to activate cell wall hydrolases, such as LytH/ActH [41], exists. Thus, it is possible in *S. aureus* the WTA machinery is positioned and regulated differently than its non-spherical counterparts in a manner that involves GpsB.

There are several other lines of evidence that give additional credence to the GpsB-WTA link. The first comes from the suppressor analysis in *L. monocytogenes* [42], where Rismondo et al. show that suppressor mutations in the WTA biosynthesis pathway can suppress the lethality of cells growing without *gpsB* at a higher non-permissive temperature. These suppressors could be freeing lipid-II/UDP-GlcNAc for the essential peptidoglycan biosynthesis pathway. It may be possible that several lipid-II and/or UDP-GlcNAc utilizing WTA proteins and peptidoglycan synthesis components could be streamlined with the help of GpsB as a central coordinator for efficient cell cycle progression.

Another line of evidence supporting a GpsB-WTA link is the connection between WTA and division site localization/selection in multiple organisms. It has been shown that WTA machinery is enriched at the division site in both *B. subtilis* and *S. aureus* from the very start of septal membrane invagination [21, 23]. Interestingly, PBP4, a likely interaction partner of GpsB [22], depends on TarO for division site localization [23].

Investigations showed that inhibition of WTA synthesis [24, 43] or prevention of WTA transfer to peptidoglycan [44–46], affect the positioning of division septum in *S. aureus*, which indicate a direct or indirect role for WTA in division site selection. Of note, at least one of the WTA ligases in *S. aureus*, SA1195 (MsrR) may interact with GpsB [22].

Additionally, division site localization of lipoteichoic acids (LTA) synthesis complex in *S. aureus* has been noted previously [31] and it was shown that multiple proteins in the LTA machinery interact with the divisome, including PBPs and the well-characterized interaction partner of GpsB, EzrA. It is important to note that LTA synthesis happens extracellularly in the periplasmic space between the high-density cell wall zone and the membrane [47]. WTA synthesis on the other hand occurs intracellularly and is exported for covalent attachment to the peptidoglycan to be part of the high-density cell wall zone [47]. It is estimated that WTA makes up nearly 60% of the cell wall composition [19].

Evidence showing the presence of septal WTA is available [19, 24, 48, 49], and multiple studies have investigated the role and importance of WTA at sites of division. It has been proposed that the maturation (such as D-alanylation) and/or accumulation of WTA may then take place subsequent to cell division in order to not interfere with (or allow) the autolysin function [19, 23, 27, 50–53]. In support of this idea, it was reported that LytF, a major autolysin in *B. subtilis*, is excluded from the lateral cell wall and localizes specifically to division sites in a WTA-dependent manner [33]. Perhaps autolysins interact with WTAs to allow for efficient separation of conjoined daughter cells in one, or a combination of, the following three ways: autolysins are actively recruited to the division sites with the aid of immature WTAs [51]; secretion of autolysins preferentially happen with WTA export at sites of division [28]; and/or septum-localized proteins such as FmtA selectively remove D-alanylation at the division site [54]. Additionally, other proteins that are translocated specifically at division sites, such as those with YSIRK signal, may rely on teichoic acids as well [55].

Taking the multiple lines of evidence into consideration, it is reasonable to postulate the presence of a multiprotein complex of complexes comprised of the divisome (including septal peptidoglycan synthesis components) and machineries involved in synthesizing both WTA and LTA at the site of cell division. As such, we propose a model taking previously published reports into account (**Fig. 7**). GpsB initially localizes to the sites of cell division in a FtsZ-dependent manner [15]. As shown in this study, TarG is preferentially enriched at the division site (**Fig. S5C**) and directly interacts with GpsB (**Fig. 5A**). Thus, we believe the TarGH WTA export complex and possibly TarO (based on its interaction with GpsB [22]; and enrichment at division site [23]) is recruited to the division site by GpsB in *S. aureus*. Subsequent to the creation of the highly crosslinked WTA-containing cell wall at the septum (shown as dark gray lines in **Fig. 7B and C**) and secretion of autolysin at the division site, regulated autolysis allows for daughter cell separation, within the scale of milliseconds [30, 56]. Immediately after cell separation, peripherally-localized GpsB [15] may facilitate the continuous incorporation of WTA along the surface of the cell to further strengthen the cell envelope. Thus, we propose that GpsB aids in the coordination of cell division and WTA synthesis/export machineries in *S. aureus*. Further investigation is necessary to shed light on the dependency of multiple crucial cellular processes on GpsB and the nature of molecular interaction between GpsB and its multiple partners. Given the significance of GpsB in multiple cellular pathways, it is an attractive drug target for the development of anti-staphylococcal/antibiotic compounds.

**Figure 7.**
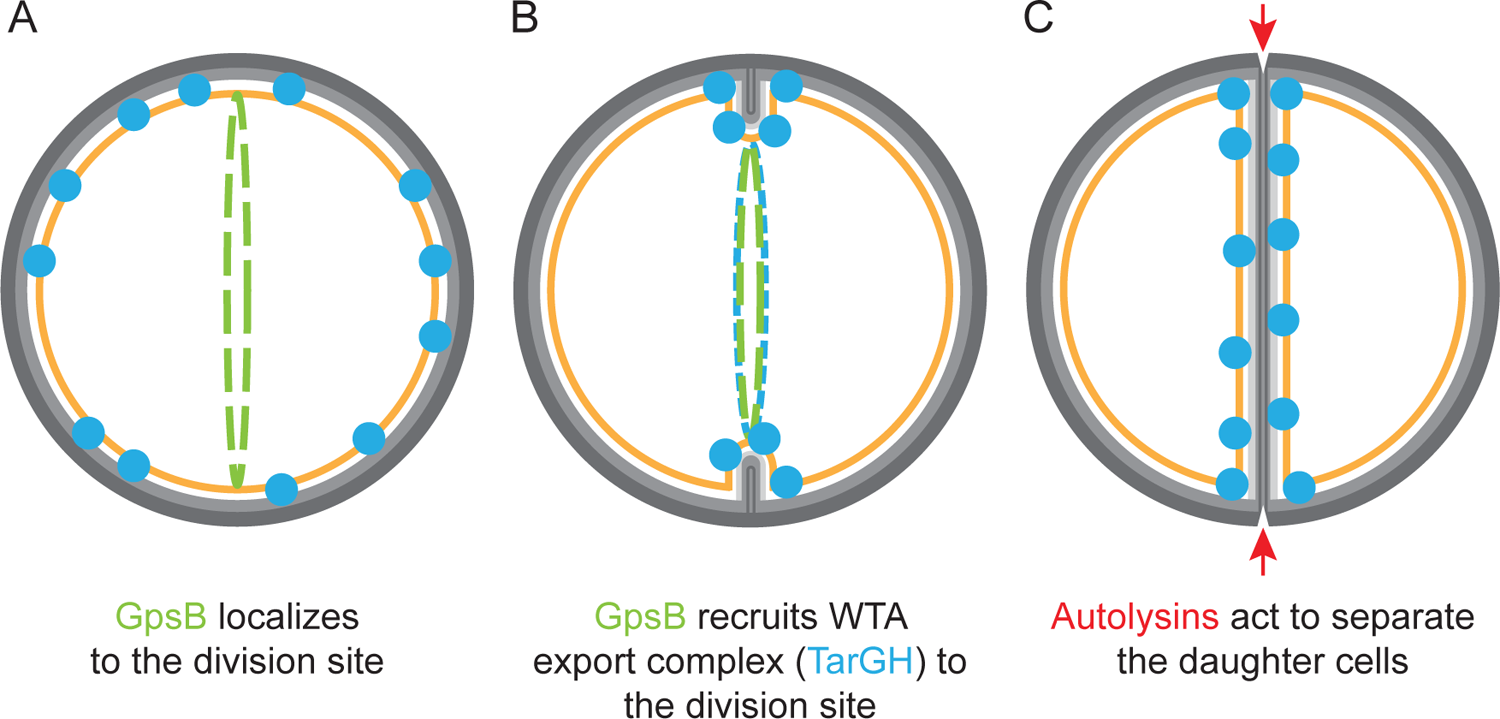
Model of GpsB-mediated coordination of cell division and wall teichoic acid export. (A) GpsB localizes to the division site in a FtsZ dependent manner [15]. (B) At the onset of membrane invagination, GpsB recruits WTA export machinery to the site of cell division. (C) Autolysins specifically act at the division septum to detach the daughter cells. GpsB – green; TarGH – blue; Autolysins – red; membrane – orange; light & dark gray – less-crosslinked cell wall and WTA-rich highly-crosslinked cell wall respectively [24, 48, 49].

## MATERIALS AND METHODS

### Suppressor Screen

Suppressor screening was carried out in the same manner as previously described [16]. Strain GG8 (*gpsB^SA^-gfp*), was plated onto LB agar plates supplemented with 1 mM IPTG to induce expression of *gpsB^SA^-gfp* and incubated at 37 °C overnight. Single colonies that were able to grow were then isolated from the plates and streaked onto new LB agar plates supplemented with 1 mM IPTG and incubated overnight at 37 °C. After confirming the ability of these strains to grow, genomic DNA was extracted and transformed into fresh wildtype PY79 cells and screened for *amyE* integration. These colonies were then streaked onto LB agar plates supplemented with 1 mM IPTG, with PY79 serving as the control, and incubated overnight at 37 °C. Strains that were not able to grow were denoted as possible extragenic mutations and sent for whole genome sequencing (MiGS Microbial Genome Sequencing center – Pittsburgh, PA). In contrast, any strains that were able to grow in the presence of the IPTG inducer were denoted as possible \**gpsB-gfp* intragenic suppressors. These strains were then screened via fluorescence microscopy to detect GFP signal and rule out possible promoter or frame shift mutations or the introduction of a premature stop codon. The genomic DNA from the \**gpsB-gfp* strains was isolated, the *amyE* locus carrying *gpsB-gfp* was PCR amplified (op36/op24), and subsequently sequenced. Analysis of the sequences was done using the ApE – A plasmid Editor (M. Wayne Davis), and multiple sequence alignments were analyzed by using the Clustal Omega multiple sequence alignment software [57].

### Strain construction

All relevant strain and oligonucleotide information is listed in Table S1. Construction of plasmids were performed with *E. coli* DH5α according to standard laboratory procedures. To generate *B. subtilis* strains carrying both mutated (labelled collectively as \**gpsB-gfp)* and unmutated copies of *gpsB-gfp* we utilized a PY79 [58] derivative that contains a second *amyE* locus (bkdB::Tn917::amyE::cat; Amy Camp). pGG4 [15] was used to clone *gpsB-gfp* into the second *amyE* locus making strain LH72. The resistance cassette was then switched from specR to ermR using pQP1 (Qi Pan) resulting in strain LH73. Following screening, genomic DNA from the \**gpsB-gfp* strains was transformed into LH73 and colonies were screened for integration at the primary *amyE* locus. This process resulted in strains LH75-LH80 that have IPTG inducible copies of both *gpsB-gfp* and \**gpsB-gfp*. Initial generation of *B. subtilis* strains containing the \**gpsB-gfp* mutations is described above (*See Suppressor Screen*). *S. aureus* strains were constructed to place \**gpsB-gfp* under the control of an IPTG inducible promoter using the pCL15 plasmid backbone [59]. The plasmid containing unmutated GpsB-GFP, pPE46 [15] was transformed into RN4220 cells resulting in strain GGS2. DNA for Y14F, ΔLEE and R72H was PCR amplified (op36/oGG2; HindIII/SalI) and cloned into the pCL15 plasmid creating plasmids pLH5-pLH9. L35S and LEErpt were made though QuikChange (Agilent) of pPE46 resulting in pPE78 and pPE80. These plasmids were then transformed into RN4220 cells creating strains LH17-20, and LH35-LH36. These plasmids were also transformed into the *S. aureus* RN4220 Δ background (SEJ1; [60]) resulting in strains LH141-LH159. The untagged Δ was similarly created through PCR amplification (op36/op38 HindIII/SphI) and cloned into the pCL15 plasmid background creating pAH1 and transformed into RN4220 cells resulting in strain AH2. Plasmids for Bacterial Two Hybrid analysis were created using pEB354 and pEB355 which carry the pUT25 and pUT18 subunits of adenylate cyclase respectively [17]. DNA for GpsB^SA^ and *GpsB^SA^ (BTH 11/BTH 12; EcoRI/XhoI) (LH39-LH40, LH43-56), TarG (BTH62/BTH63; EcoRI/Xho1) (SKB1-SKB2), and TarH (BTH 60/BTH61; EcoRI/Xho1) (SKB3-SKB4), was PCR amplified and cloned into both pEB354 and pEB355. To create the gfp-tagH strain in *B. subtilis* cells, chromosomal DNA from Bacillus Genetic Stock Center (BGSC) 1A1119 [21] was transformed into PY79 cells to create PE528. Similarly, to create the CRISPRi TagG/TagH strains, chromosomal DNA was extracted from strains BEC35710 and BEC35700 (from BGSC) respectively and transformed into PY79 cells to make SK15-SK16. Then, to create cells that contained the CRISPRi knockdown for TagG or TagH as well as IPTG inducible *gpsB^SA^-gfp*, the same DNA was transformed into LH72 resulting in SK17-SK18. Finally, to create *gfp-tarG*, DNA was PCR amplified (oLH11/12; SalI/BamHI and oLH13/14; BamHI/EcoRI) and ligated into the pJB67 vector [61] resulting in pLH64 which was then transformed into RN4220 cells to make LH136.

### Spot Titer Assay

The spot titer assays for *B. subtilis* strains were carried out on LB agar plates supplemented with 1 mM IPTG where needed to induce the expression of *gpsB*^SA^*-gfp* or \**gpsB*^SA^*-gfp*. Cultures of the strains were first grown to mid-logarithmic phase (OD_600_=0.4-0.6) and subsequently standardized to OD_600_=0.1. The standardized cultures were then serially diluted and 1 µl of the liquid culture was spotted onto the appropriate plate. These plates were then incubated overnight at 37 °C. Spot titer assays for *S. aureus* strains were completed in the same manner using Tryptic Soy Agar plates supplemented with 10 µg/ml chloramphenicol, and where needed, 1 mM IPTG. The serial dilutions of the standardized cultures were spotted onto the plates with a volume of 1 µl.

### Fluorescence Microscopy

Florescence microscopy was carried out as previously described [62]. Overnight cultures of *B. subtilis* strains in LB liquid media, or *S. aureus* strains in TSB supplemented with 10 µg/ml chloramphenicol (pCL15 backbone) or 5 µg/ml erythromycin (pJB67 backbone), were diluted to OD_600_=0.1 and grown at 37 °C to mid logarithmic phase (OD_600_= 0.4-0.6). Then, for *B. subtilis* cultures, where needed, 1 mM IPTG or 1% xylose was added to the cultures and grown for an additional 3 h. For *S. aureus*, 1 mM IPTG (pCL15) or 1.25 µM CdCl_2_ (pJB67) was added to the cultures (except GFP-TarG where no inducer was added) and where needed, 20 μg/mL targocil was also added. Cells were then grown for an additional 1 h. Following incubation, 1 ml aliquots were then pelleted and resuspended g/ml SynaptoRed fluorescent dye to allow for visualization of the μl) was then transferred to a glass bottom dish (Mattek) and covered with a 1% agarose pad. Samples were imaged on a DeltaVision Core microscope system (Applied Precision/GE Healthcare) equipped with a Photometrics CoolSnap HQ2 camera and an environmental chamber. Seventeen planes were acquired every 200 nm and the data were deconvolved using SoftWorx software. Cell diameter was measured using ImageJ and analysis completed in GraphPad Prism 9.

### Bacterial Two Hybrid Assay

Plasmids carrying our genes of interest were transformed into *E. coli* BTH101 cells and μg/ml ampicillin and 50 μ incubated at 30 °C for 48 h. A positive control strain was made by transforming PE87 and PE88 carrying pUT25-zip and pUT18-zip into the BTH101 cells. A negative control was made by transforming empty pEB354 and pEB355 into the cells. The resulting colonies harboring the pairs of plasmids of interest were then isolated and grown overnight in liquid LB media at 30 °C supplemented with 100 μg/ml ampicillin, 50 μg/ml kanamycin, and 0.5 mM IPTG. The following day, 2 µl of culture was spotted onto MacConkey plates (BD Biosciences) supplemented with 10 mg/ml maltose, 100 μg/ml ampicillin, 50 μg/ml kanamycin, and 0.5 mM IPTG. Plates were then incubated for 24 h β-galactosidase assays were completed in the 96 well plate reader as previously described with some modification [63]. A mixture of 450 µl Z buffer, 120 µl β-mercaptoethanol, 285 µl (polymyxin B 20 mg/ml), and 60 µl of culture for each strain tested was prepared. 200 µl of each mixture was then transferred in triplicate to a 96 well plate. Readings were taken in a BioTek plate reader and Miller units calculated. Results were graphed using GraphPad Prism 9.

### Immunoblotting

Cells were grown overnight and the next morning diluted to an OD_600_=0.1. At OD_600_= 0.4-0.6, IPTG was added to the appropriate cultures and then grown for an additional 1 h (*S. aureus*) or 3 h (*B. subtilis*). Aliquots of cultures (1 ml) were then collected and pelleted. Cell lysis of *B. subtilis* cells was completed by using a protoplast buffer containing 0.5 M sucrose, 20 mM MgCl_2_, 10 mM KH_2_PO_4_, and 0.1 mg/ml lysozyme. *S. aureus* cells (in RN4220 Δ*spa* background [60]) were resuspended in 500 µl PBS with 5 µl lysostaphin (1 mg/ml in 20 mM sodium acetate) and incubated for 30 min at 37 °C. 1 µl of DNAse A (1U/µl) was then added and allowed to incubate for an additional 30 min. Samples were then prepared for SDS-PAGE analysis. After electrophoresis, the samples were transferred to a membrane and probed with rabbit antisera raised against GFP (K. Ramamurthi) and SigA (M. Fujita) for *B. subtilis*, and GpsB-GFP for *S. aureus* with total protein visualized using the GelCode Blue Safe Protein Stain (ThermoFisher).

### WTA Extraction

WTAs were extracted and visualized as previously described [64]. Briefly, cultures of PES5 and PES13 were grown overnight and back diluted to an OD_600_ of 0.1 the next day. Cultures were then grown to mid-log before 1 mM IPTG was added, and then the cultures were grown for an additional 3 h. Cells were standardized to the same OD (OD_600_ of 1) and then pelleted and washed in SDS buffer. Cells were then boiled for 1 h and subsequently extensively washed in SDS buffer. The cells were then subjected to proteinase K treatment, and following washes in sterile distilled water, the WTAs were extracted with NaOH overnight. The following day the tubes were centrifuged to separate the WTA from leftover debris. The supernatant was then run on a Native PAGE gel and visualized with Alcian Blue (1:20 dilution of 1.25% stock solution in 2% acetic acid) followed by Silver Staining (ThermoFisher) following manufacturer protocols and imaged on the Bio-Rad Chemidoc MP Imaging System.

### Autolysis Assay

Autolysis assays were carried out as previously described [28]. Overnight cultures were back diluted to an OD_600_ of 0.1. Cultures were grown at 37°C to mid-log (OD_600_ 0.4-0.6) and then 1mM IPTG was added and when needed targocil (5 µg/ml). Cells were then grown for an additional hour. Cells were then pelleted, washed twice in cold H_2_O, and standardized to an OD_600_ of 0.8. Cells were then spun and resuspended in 0.5M Tris-HCl (pH 7.2) and 0.05% Triton -X 100. OD_600_ was monitored in a 96-well plate reader for 10 hours at 37°C. Results were graphed using Graphpad Prism 9.

### Structural analysis and Multiple Sequence Alignment

The 3-dimensional model for the N-terminal domain of GpsB^Sa^ (res. 1-70) was generated with the SWISS-MODEL homology-model server [18]. GpsB suppressor mutants, and simulated interactions with homologous PBP complexes were generated with PyMOL (Schrödinger, LLC). PDB coordinates for *S. pneumoniae*, *B. subtilis*, and *L. monocytogenes* GpsB in complex with their associated PBPs were retrieved from the PDB, with accession IDs 6GQN, 6GP7, and 6GPZ [14].

## ACKNOWLEDGEMENTS

We thank our lab members for comments on the manuscript. This work was funded by the National Institutes of Health grants (R35GM133617; PE) and (R21AI164775; YC and PE).

## AUTHOR CONTRIBUTIONS

The conception and design of the study (LH, CS, PE), data acquisition (LH, SK, MS, AH, CS), analysis and/or interpretation of the data (LH, SK, MS, CS, YC, PE), and writing of the manuscript (LH, MS, YC, PE).

