## Supplemental Information for "GpsB coordinates cell division and cell surface decoration by wall teichoic acids in *Staphylococcus aureus*"

**Supplemental Table and Figure Legends**

Figure S1

(A) Stable production of *GpsB$^{SA}$-GFP was confirmed with western blot analysis of PY79 *B. subtilis* cells expressing *gpsB$^{SA}$-gfp* (GG8) or **gpsB$^{SA}$-gfp* (CS89-CS93, PE377, PE448) both with and without the addition of 1 mM IPTG. (B) The L35S *GpsB-GFP (PE448) mutation shows a clevage product detected via western blot. GpsB$^{SA}$-GFP (GG8) and free GFP (RL4709) are shown for comparison. Blots probed with anti-GFP antibodies and anti-sigA for a loading control. (C) Localization of GpsB$^{SA}$-GFP (i) (GG8) and *GpsB$^{SA}$-GFP (ii – viii) (CS89-CS93, PE377, PE448) in *B. subtilis* PY79 cells. Images taken 3 h after the addition of 1 mM IPTG and membranes were visualized with SynaptoRed membrane dye. Scale bar is 1 μm.

Figure S2

Investigation of *GpsB dominant negative phenotypes. (A) Spot titer assay with *B. subtilis* PY79 cells co-expressing *gpsB$^{SA}$-gfp* (LH73) and one of the **gpsB$^{SA}$-gfp* intragenic suppressor mutations (LH75-LH80). Cultures were serially diluted and spotted onto plates in the absence (left) and presence (right) of 1 mM IPTG. (B) BACTH assay testing pairwise interactions of *GpsB$^{SA}$ (LH43, LH45, LH47, LH49, LH51, LH53, LH55) with WT GpsB$^{SA}$ (LH40). Interactions on solid media with a dark pink color indicating a positive interaction. Image taken after 48 h of incubation (top). β galactosidase assay testing the same pairwise interactions. Bars show calculated Miller units with the dashed line marking the cutoff from the negative control (bottom). (C) Spot titer assay of RN4220 *S. aureus* cells harboring *GpsB$^{SA}$-GFP mutations (PE355, GGS2, LH17-LH20, LH32, LH35, LH36)  and plated on media containing 1 mM IPTG. (D) Production of GpsB$^{SA}$-GFP (LH141) and *GpsB$^{SA}$-GFP (LH142-LH144, LH159-LH162) in *S.*

*aureus* RN4220 Δ*spa* cells. Blots were probed with antibody raised against GpsB$^{SA}$-GFP and a total protein stained gel was used as a loading control.

Figure S3

Superimposition of the crystal structures of *S. pneumoniae (*PDB ID 6GQA), *B. subtilis* (PDB ID 4UG3), *L. monocytogenes* (PDB ID 4UG1), GpsB and the *S. aureus* homology model. (A) Side view. (B) Top view. (C) The equivalent of *S. aureus* D41 is strictly conserved in *S. pneumoniae, B. subtilis, and L. monocytogenes*, where it functions as a hydrogen bond acceptor for three residues. These interactions are an important force that stabilizes this region. The subscripts A and B denote the two protomers in the dimer.

Figure S4

Homology model of *S. aureus* GpsB. The N-terminal LEELR section of α-helix 2 with (a) native and (b) inserted LEE residues colored in yellow.

Figure S5

(A) β galactosidase assay testing pairwise interactions of GpsB$^{SA}$ (LH39-LH40) with TarG (SKB1-SKB2) and TarH (SKB3-SKB4). Bars show calculated Miller units with the dashed line marking the cutoff from the negative control. (B) Purified WTA extracts from strains of SH1000 *S. aureus* cells containing the empty vector (control) (PES5) or a plasmid for GpsB$^{SA}$ overexpression (PES13). Cells were grown to mid-log phase and then induced with 1 mM IPTG for 3 h prior before collection and extraction of WTAs. Extracts were analyzed via electrophoresis on a native PAGE gel and visualized through Alcian blue and silver staining. (C)

Fluorescence micrographs showing localization of GFP-TarG in RN4220 *S. aureus* cells (LH136) imaged at mid-log phase with no inducer added. Scale bar 1 μm. (D) Measurement of autolysis in strains overproducing GpsB$^{SA}$ (GGS1) and GpsB$^{\Delta LEE}$ (AH2) and an empty vector control (PE355) both with and without treatment of 5 μg/ml targocil.

Table S1

Strains and oligonucleotides used in this study.

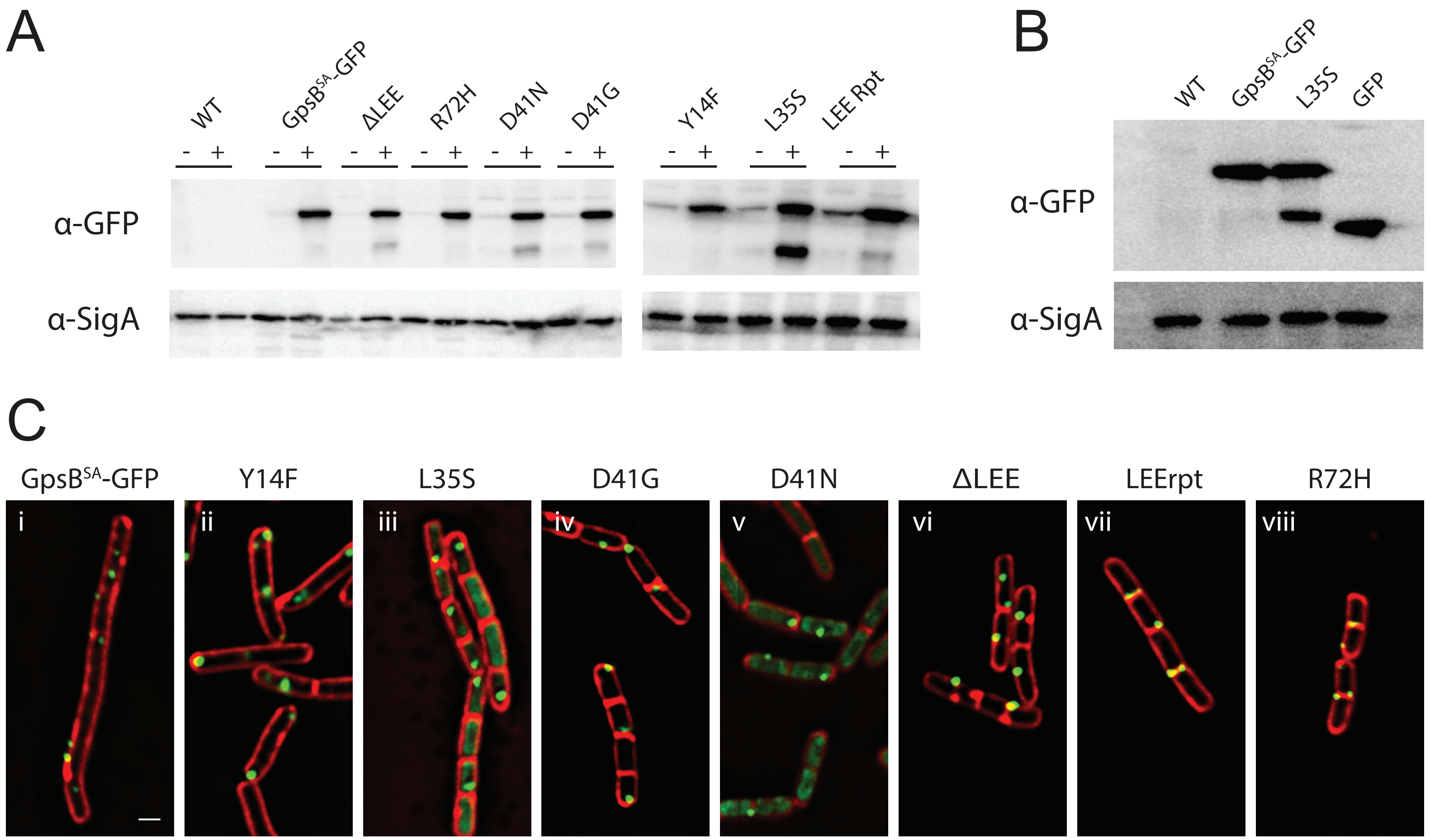
A
WT
GpsB^SA-GFP
ΔLEE
R72H
D41N
D41G
Y14F
L35S
LEE Rpt
α-GFP
α-SigA
B
WT
GpsB^SA-GFP
L35S
GFP
α-GFP
α-SigA
C
GpsB^SA-GFP
Y14F
L35S
D41G
D41N
ΔLEE
LEErpt
R72H
i
ii
iii
iv
v
vi
vii
viii

Figure S1

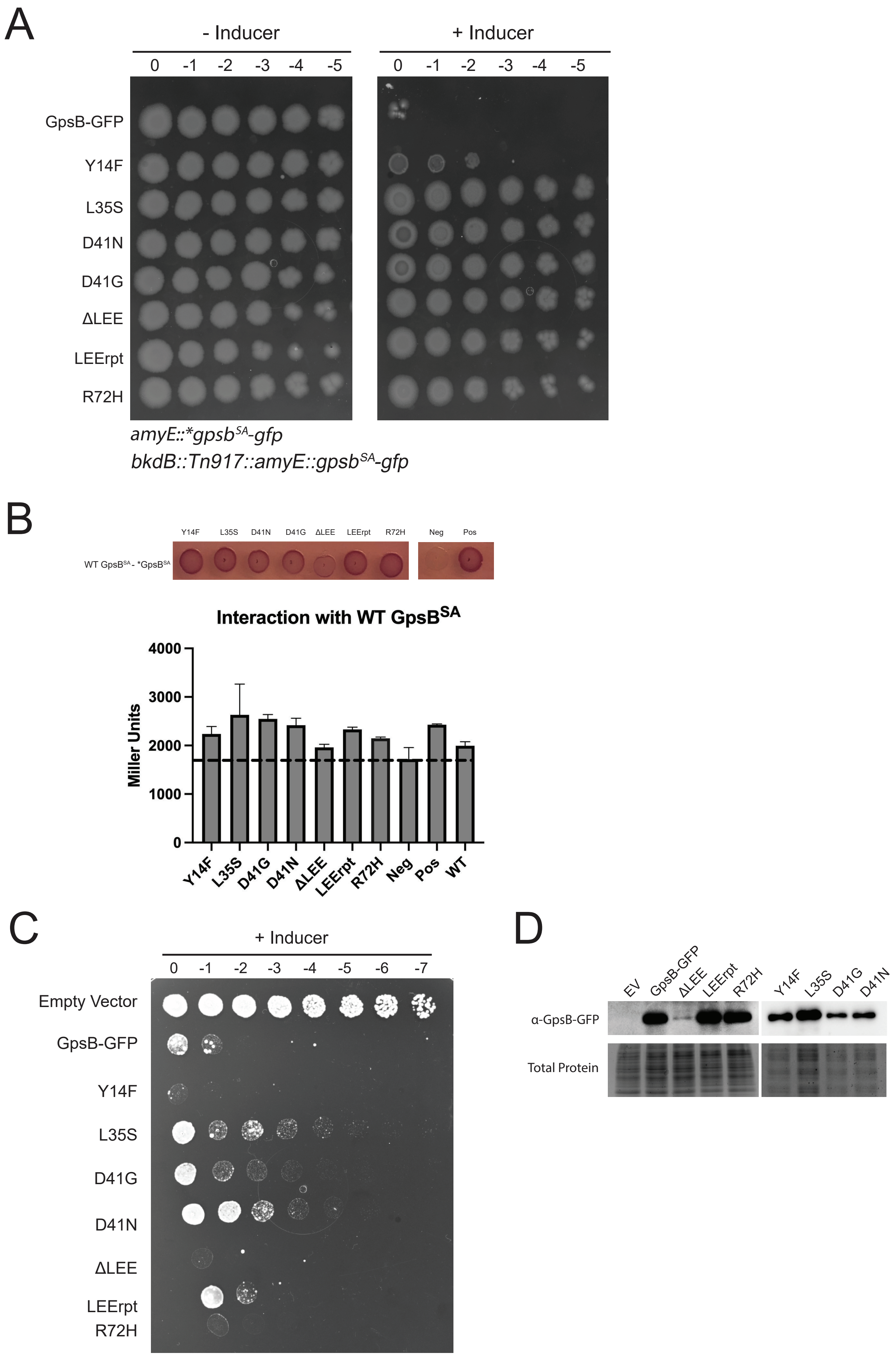
A
- Inducer
+ Inducer
0 -1 -2 -3 -4 -5
0 -1 -2 -3 -4 -5
GpsB-GFP
Y14F
L35S
D41N
D41G
ΔLEE
LEErpt
R72H
*amyE::*gpsb$^{SA}$-gfp*
*bkdB::Tn917::amyE::gpsb$^{SA}$-gfp*
B
Y14F L35S D41N D41G ΔLEE LEErpt R72H Neg Pos
WT GpsB$^{SA}$ - *GpsB$^{SA}$
Interaction with WT GpsB$^{SA}$
Miller Units
0
1000
2000
3000
4000
Y14F L35S D41G D41N ΔLEE LEErpt R72H Neg Pos WT
C
+ Inducer
0 -1 -2 -3 -4 -5 -6 -7
Empty Vector
GpsB-GFP
Y14F
L35S
D41G
D41N
ΔLEE
LEErpt
R72H
D
EV GpsB-GFP ΔLEE LEErpt R72H Y14F L35S D41G D41N
α-GpsB-GFP
Total Protein

Figure S2

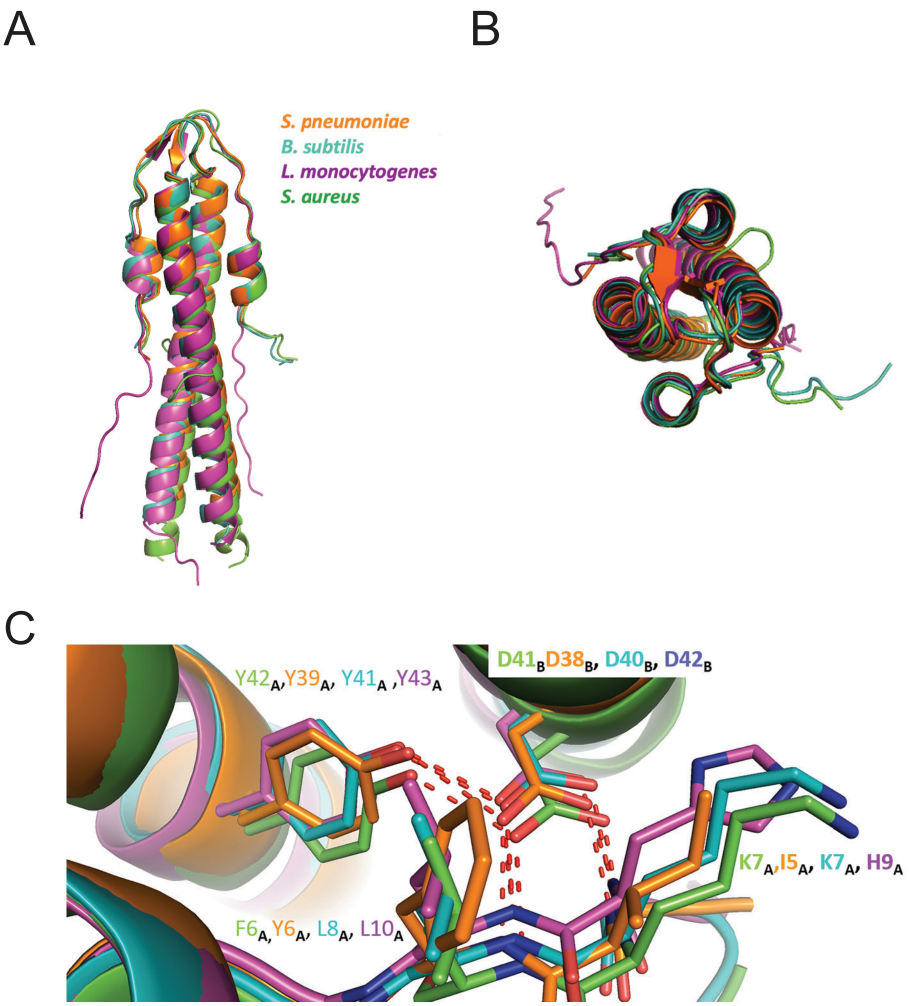
A
B
C
S. pneumoniae
B. subtilis
L. monocytogenes
S. aureus
Y42A,Y39A, Y41A ,Y43A
D41BD38B, D40B, D42B
K7A,I5A, K7A, H9A
F6A,Y6A, L8A, L10A

Figure S3

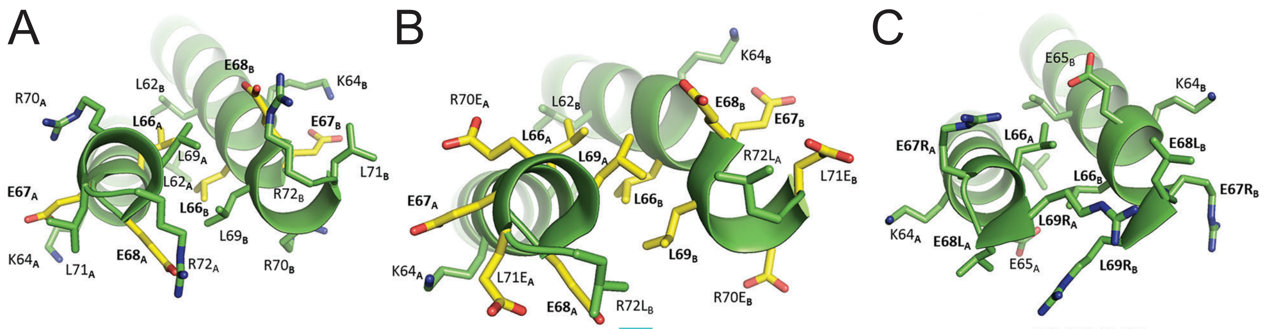

66-LEEELR-72

LEE

66-LEELEEL-72

LEE repeat

66-LRLR-69

LEE deletion

Figure S4

A
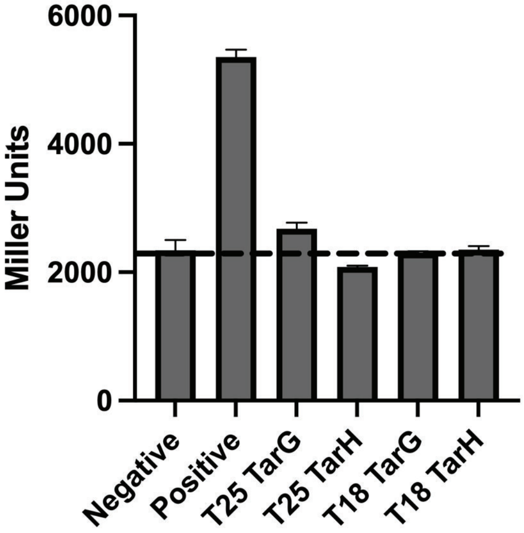

B
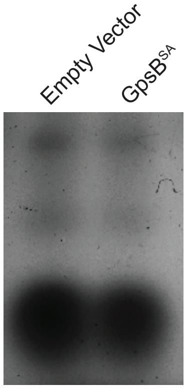

C
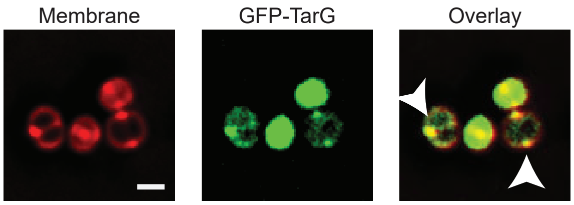

D
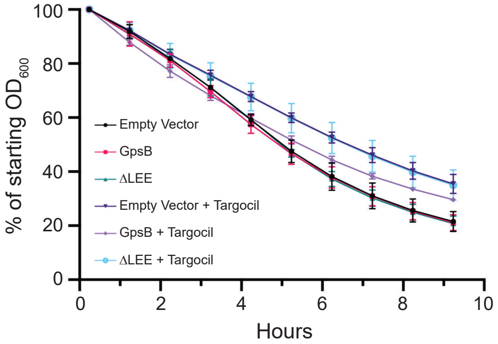

Figure S5

**Table S1** Strains and oligonucleotides used in this study.

### Strains used in this study

| Species | Strain | Genotype | Source |
|---|---|---|---|
| *B. subtilis* | PY79 | Wildtype | [1] |
| *B. subtilis* | GG8 | amyE::$P_{hyperspank}$-gpsB$^{SA}$-gfp | [2] |
| *B. subtilis* | PE448 | amyE::$P_{hyperspank}$-*gpsB$^{SA-L35S}$-gfp | [2] |
| *B. subtilis* | PE377 | amyE:: $P_{hyperspank}$-*gpsB$^{SA-LEErpt}$-gfp | This study |
| *B. subtilis* | CS89 | amyE:: $P_{hyperspank}$-*gpsB$^{SA-D41N}$ -gfp | This study |
| *B. subtilis* | CS90 | amyE:: $P_{hyperspank}$-*gpsB$^{SA-\Delta LEE}$ -gfp | This study |
| *B. subtilis* | CS91 | amyE:: $P_{hyperspank}$-*gpsB$^{SA-R72H}$ -gfp | This study |
| *B. subtilis* | CS92 | amyE:: $P_{hyperspank}$-*gpsB$^{SA-Y14F}$ -gfp | This study |
| *B. subtilis* | CS93 | amyE:: $P_{hyperspank}$-*gpsB$^{SA-D41G}$ -gfp | This study |
| *B. subtilis* | LH72 | bkdB::Tn917::amyE::$P_{hyperspank}$-gpsB-gfp (specR) | This study |
| *B. subtilis* | LH73 | bkdB::Tn917::amyE::$P_{hyperspank}$-gpsB-gfp (ermR) | This study |
| *B. subtilis* | LH75 | bkdB::Tn917::amyE::$P_{hyperspank}$-gpsB-gfp<br>amyE::$P_{hyperspank}$-*gpsB$^{SA-L35S}$-gfp | This study |
| *B. subtilis* | LH79 | bkdB::Tn917::amyE::$P_{hyperspank}$-gpsB-gfp<br>amyE:: $P_{hyperspank}$-*gpsB$^{SA-LEErpt}$-gfp | This study |
| *B. subtilis* | LH76 | bkdB::Tn917::amyE::$P_{hyperspank}$-gpsB-gfp<br>amyE:: $P_{hyperspank}$-*gpsB$^{SA-D41N}$ -gfp | This study |
| *B. subtilis* | LH78 | bkdB::Tn917::amyE::$P_{hyperspank}$-gpsB-gfp<br>amyE:: $P_{hyperspank}$-*gpsB$^{SA-\Delta LEE}$ -gfp | This study |
| *B. subtilis* | LH80 | bkdB::Tn917::amyE::$P_{hyperspank}$-gpsB-gfp<br>amyE:: $P_{hyperspank}$-*gpsB$^{SA-R72H}$ -gfp | This study |
| *B. subtilis* | LH74 | bkdB::Tn917::amyE::$P_{hyperspank}$-gpsB-gfp<br>amyE:: $P_{hyperspank}$-*gpsB$^{SA-Y14F}$ -gfp | This study |
| *B. subtilis* | LH77 | bkdB::Tn917::amyE::$P_{hyperspank}$-gpsB-gfp<br>amyE:: $P_{hyperspank}$-*gpsB$^{SA-D41G}$ -gfp | This study |
| *B. subtilis* | RL4709 | amyE::$P_{hyperspank}$-gfp | (R. Losick Lab) |
| *B. subtilis* | SK15 | lacA::Pxyl-dcas9 amyE::Pveg-sgRNA (tagG) | This study; derived from BEC35710 (^BGSC) |
| *B. subtilis* | SK16 | lacA::Pxyl-dcas9 amyE::Pveg-sgRNA (tagH) | This study; derived from BEC35700 (^BGSC) |
| *B. subtilis* | SK17 | lacA::Pxyl-dcas9 amyE::Pveg-sgRNA (tagG);<br>bkdB::Tn917::amyE::$P_{hyperspank}$-gpsB-gfp | This study |
| *B. subtilis* | SK18 | lacA::Pxyl-dcas9 amyE::Pveg-sgRNA (tagH);<br>bkdB::Tn917::amyE::$P_{hyperspank}$-gpsB-gfp | This study |
| *B. subtilis* | PE528 | tagH::$P_{xyl}$-gfp-tagH (1-648) | This study; derived from [3] |
| *B. subtilis* | GG19 | amyE::$P_{hyperspank}$-gpsB$^{BS}$-gfp | [2] |
| *S. aureus* | SH1000 | Wildtype | [4] |
| *S. aureus* | RN4220 | Wildtype | Lab stock |
| *S. aureus* | SEJ1 | RN4220 Δ*spa* | [5] |
| *S. aureus* | PES5 | SH1000 with pCL15 Empty Vector | [2] |
| *S. aureus* | PE355 | RN4220 with pCL15 Empty Vector | [2] |
| *S. aureus* | PES6 | SH1000 pCL15 backbone $P_{spac}$-gpsB$^{SA}$-gfp | [2] |
| *S. aureus* | GGS2 | RN4220 pCL15 backbone $P_{spac}$-gpsB$^{SA}$-gfp | [2] |
| *S. aureus* | PES13 | SH1000 pCL15 backbone $P_{spac}$-gpsB$^{SA}$ | [2] |
| *S. aureus* | GGS1 | RN4220 pCL15 backbone $P_{spac}$-gpsB$^{SA}$ | [2] |
| *S. aureus* | LH36 | RN4220 pCL15 backbone $P_{spac}$-*gpsB$^{SA\ L35S}$-gfp | [2] |
| *S. aureus* | LH35 | RN4220 pCL15 backbone $P_{spac}$-*gpsB$^{SA\ LEErpt}$-gfp | This study |
| *S. aureus* | LH19 | RN4220 pCL15 backbone $P_{spac}$-*gpsB$^{SA\ D41N}$-gfp | This study |
| *S. aureus* | LH17 | RN4220 pCL15 backbone $P_{spac}$-*gpsB$^{SA\ \Delta LEE}$-gfp | This study |
| *S. aureus* | LH18 | RN4220 pCL15 backbone $P_{spac}$-*gpsB$^{SA\ R72H}$-gfp | This study |
| *S. aureus* | LH32 | RN4220 pCL15 backbone $P_{spac}$-*gpsB$^{SA\ Y14F}$-gfp | This study |
| *S. aureus* | LH20 | RN4220 pCL15 backbone $P_{spac}$-*gpsB$^{SA\ D41G}$-gfp | This study |
| *S. aureus* | LH136 | RN4220 pJB67 backbone $P_{cad}$-gfp-tarG | This study |
| *S. aureus* | AH2 | RN4220 pCL15 backbone $P_{spac}$-*gpsB$^{SA\ \Delta LEE}$ | This study |
| *S. aureus* | LH140 | RN4220 with pCL15 Empty Vector | This study |
| *S. aureus* | LH141 | RN4220 Δspa pCL15 backbone $P_{spac}$-gpsB$^{SA}$-gfp | This study |
| *S. aureus* | LH162 | RN4220 Δspa pCL15 backbone $P_{spac}$-*gpsB$^{SA\ L35S}$-gfp | This study |
| *S. aureus* | LH144 | RN4220 Δspa pCL15 backbone $P_{spac}$-*gpsB$^{SA\ LEErpt}$-gfp | This study |
| *S. aureus* | LH160 | RN4220 Δspa pCL15 backbone $P_{spac}$-*gpsB$^{SA\ D41N}$-gfp | This study |
| *S. aureus* | LH142 | RN4220 Δspa pCL15 backbone $P_{spac}$-*gpsB$^{SA\ \Delta LEE}$-gfp | This study |

| | | | |
|---|---|---|---|
| *S. aureus* | LH143 | RN4220 ∆spa pCL15 backbone $P_{spac}$-*$gpsB^{SA\ R72H}$-*gfp* | This study |
| *S. aureus* | LH161 | RN4220 ∆spa pCL15 backbone $P_{spac}$-*$gpsB^{SA\ Y14F}$-*gfp* | This study |
| *S. aureus* | LH159 | RN4220 ∆spa pCL15 backbone $P_{spac}$-*$gpsB^{SA\ D41G}$-*gfp* | This study |
| *E. coli* | BTH101 | *Adenylate cyclase deficient reporter strain for BACTH; F', cya-99, araD139, galE15, galK16, rpsL1 ($Str^R$), hsdR2, mcrA1, mcrB1, relA1* | [6] |
| *E. coli* | LH39 | T18-linker $gpsB^{SA}$ | This study |
| *E. coli* | LH40 | T25-linker $gpsB^{SA}$ | This study |
| *E. coli* | LH43 | T18-linker *$gpsB^{SA\ LEErpt}$ | This study |
| *E. coli* | LH44 | T25-linker *$gpsB^{SA\ LEErpt}$ | This study |
| *E. coli* | LH45 | T18-linker *$gpsB^{SA\ L35S}$ | This study |
| *E. coli* | LH46 | T25-linker *$gpsB^{SA\ L35S}$ | This study |
| *E. coli* | LH47 | T18-linker *$gpsB^{SA\ \Delta LEE}$ | This study |
| *E. coli* | LH48 | T25-linker *$gpsB^{SA\ \Delta LEE}$ | This study |
| *E. coli* | LH49 | T18-linker *$gpsB^{SA\ Y14F}$ | This study |
| *E. coli* | LH50 | T25-linker *$gpsB^{SA\ Y14F}$ | This study |
| *E. coli* | LH51 | T18-linker *$gpsB^{SA\ D41G}$ | This study |
| *E. coli* | LH52 | T25-linker *$gpsB^{SA\ D41G}$ | This study |
| *E. coli* | LH53 | T18-linker *$gpsB^{SA\ D41N}$ | This study |
| *E. coli* | LH54 | T25-linker *$gpsB^{SA\ D41N}$ | This study |
| *E. coli* | LH55 | T18-linker *$gpsB^{SA\ R72H}$ | This study |
| *E. coli* | LH56 | T25-linker *$gpsB^{SA\ R72H}$ | This study |
| *E. coli* | SKB3 | T18-linker *tarH* | This study |
| *E. coli* | SKB4 | T25-linker *tarH* | This study |
| *E. coli* | SKB1 | T18-linker *tarG* | This study |
| *E. coli* | SKB2 | T25-linker *tarG* | This study |
| *E. coli* | PE87 | pUT25-zip | [6] |
| *E. coli* | PE88 | pUT18-zip | [6] |
| *E. coli* | PE84 | pEB355 | [6] |
| *E. coli* | PE83 | pEB354 | [6] |
| *E. coli* | pCL15 | pCL15 | [7] |
| *E. coli* | pLH5 | pCL15 backbone $P_{spac}$-*$gpsB^{SA\ \Delta LEE}$-*gfp* | This study |
| *E. coli* | pLH6 | pCL15 backbone $P_{spac}$-*$gpsB^{SA\ R72H}$-*gfp* | This study |
| *E. coli* | pLH7 | pCL15 backbone $P_{spac}$-*$gpsB^{SA\ D41N}$-*gfp* | This study |
| *E. coli* | pLH8 | pCL15 backbone $P_{spac}$-*$gpsB^{SA\ D41G}$-*gfp* | This study |
| *E. coli* | pLH13 | pCL15 backbone $P_{spac}$-*$gpsB^{SA\ Y14F}$-*gfp* | This study |
| *E. coli* | pPE78 | pCL15 backbone $P_{spac}$-*$gpsB^{SA\ LEErpt}$-*gfp* | This study |
| *E. coli* | pPE80 | pCL15 backbone $P_{spac}$-*$gpsB^{SA\ L35S}$-*gfp* | This study |
| *E. coli* | pPE46 | pCL15 backbone $P_{spac}$-$gpsB^{SA}$-*gfp* | [2] |
| *E. coli* | pPE45 | pCL15 backbone $P_{spac}$-$gpsB^{SA}$ | [2] |
| *E. coli* | pLH64 | pJB67 backbone $P_{cad}$-*gfp-tarG* | This study |
| *E. coli* | pAH1 | pCL15 backbone $P_{spac}$-*$gpsB^{SA\ \Delta LEE}$ | This study |

*denotes mutant $gpsB^{SA}$

^BGSC - Bacillus Genetic Stock Center

### Oligonucleotides used in this study

| Primer | Sequence (5' to 3') |
|---|---|
| oP36 | AAAAAGCTTACATAAGGAGGAACTACTATGTCAGATGTTTCATTGAAATTATCAGCA |
| oP38 | AAAGCATGCTTATTTACCAAATACAGCTTTTTCTAAGTTTGA |
| oGG2 | AAAGTCGACTTATTTGTATAGTTCATCCATGCC |
| BTH11 | AATAAGAATTCATGTCAGATGTTTCATTGAAATTATCAGC |
| BTH12 | GCTGTATTTGGTAAATAACTCGAGTTATT |
| BTH61 | AATAAGAATTCATGAACGTTTCGGTAAACATTAAAAATG |
| BTH62 | AATAACTCGAGTTATTTAATAACGAAGCGGGACTCATCG |
| BTH63 | AATAAGAATTCATGTCAGCAATAGGAACAG |
| BTH64 | AATAACTCGAGTTACAAGAAGTCTGCAAATTGATCTC |
| oLH11 | AATAAGTCGACACATAAGGAGGAACTACTATGAGTAAAGGAGAAGAACTTTTCAC |
| oLH12 | AATAAGGATCCTTTGTATAGTTCATCCATGCC |
| oLH13 | AATAAGGATCCATGTCAGCAATAGGAACAG |
| oLH14 | AATAAGAATTCTTACAAGAAGTCTGCAAATTG |
